# Integrative spatial multi-omics reveals prognostic tumor niches in female genital tumors

**DOI:** 10.1101/2025.11.24.690313

**Authors:** Tian Xu, Siyu Lin, Yuling Wu, Yufeng He, Zongxu Zhang, Yanping Zhao, Peng Zhang, Luyang Zhao, Wen Yang, Mingxia Ye, Rui Zhang, Liang Wen, Pengfei Ren, Ce Luo, Zhe Zhang, Zexian Zeng, Yuanguang Meng

## Abstract

Female genital tumors (FGTs), including ovarian, endometrial, and cervical cancers, pose a major global health challenge, yet their spatial and molecular determinants of progression and therapy resistance remain poorly understood. Here we present a large-scale integrative spatial and multi-omics atlas of FGTs, combining exome, transcriptome, methylome, and miRNA sequencing with high-resolution Visium HD spatial transcriptomics, profiling over 62 million cells from 100 tumors. We show that HPV-driven suppression of TGFβ signaling in cervical cancer disrupts myofibroblast (myCAF) formation and reshapes immune infiltration, while serous tumors exhibit extensive oncogene amplifications that activate MYC and PI3K pathways, linking chromosomal instability (CIN) to proliferative, immune-cold phenotypes. In high-grade serous ovarian cancer, we identify a previously unrecognized malignant domain, the Proliferating Core Desert (PCD) niche, defined by CIN-high epithelial enrichment, immune exclusion, stromal protection, and chemotherapy resistance, maintained through crosstalk with macrophages and fibroblasts. Finally, we develop POWER, a pathology foundation model that predicts PCD niche abundance from routine H&E slides, enabling robust stratification of prognosis and anti-angiogenic therapy response.

## Introduction

Female genital tumors (FGTs) include a heterogeneous group of malignancies originating from the ovary, fallopian tube, corpus uteri, cervix uteri, vagina, and vulva. Collectively, these cancers account for ∼15% of newly diagnosed malignancies and cancer-related mortality among females globally^1^, underscoring their significant impact on women’s health. Despite their prevalence, FGTs remain clinically challenging due to their diverse etiologies, complex tumor microenvironment (TME), heterogeneous clinical courses, and variable responses to therapy^2,3^. Dissecting how intratumoral spatial heterogeneity contributes to therapy resistance is therefore critical for biomarker discovery and therapeutic innovation.

Although tumors arise from distinct anatomical sites, increasing evidence suggests that cancers originating from related tissues often share genetic drivers and treatment susceptibilities^4,5^. FGTs, in particular, arise from a common embryonic origin in the Müllerian ducts and display overlapping histopathological features^6,7^. Previous studies have identified shared genomic alterations between ovarian cancer (OC) and endometrial cancer (EC)^8,9^, including similarities in serous and endometrioid subtypes. Serous tumors across both OC and EC are characterized by frequent TP53 mutations, extensive chromosomal instability (CIN), and widespread copy-number variation (CNV), whereas endometrioid tumors typically lack TP53 mutations and exhibit minimal CNV^10,11^. These subtype-specific genomic patterns underlie distinct biological behaviors, therapeutic responses, and patient outcomes, highlighting the need for a unified, pan-gynecologic analysis to capture both shared and divergent features of FGTs.

Among these, CIN has emerged as a defining feature of high-grade serous ovarian cancer (HGSOC)^12^ the most aggressive and lethal subtype of OC, accounting for a disproportionate share of mortality across FGTs^13^. CIN has been linked to tumor progression, metastasis, therapeutic resistance, and extensive intratumoral heterogeneity^14–16^. Importantly, CNV burden has also been associated with reduced immune infiltration^17,18^, providing a potential explanation for the “immune cold” phenotype observed in most HGSOC tumors and their poor response to immune checkpoint blockade^19^. However, much of this knowledge is derived from bulk-level genomics and transcriptomics, or from single-cell RNA sequencing, which lack the spatial resolution required to map intratumoral CIN heterogeneity alongside immune infiltration patterns. Thus, integrative approaches combining high-resolution spatial transcriptomics with complementary multi-omics profiling are essential to resolve cancer cell states in tissue context and TME interactions.

Here, we present a high-resolution spatial atlas of FGTs generated using Visium HD spatial transcriptomics^20^, profiling over 62 million cells across 100 tumor sections, including 58 OC, 19 EC, and 23 cervical cancer (CC) samples. We also generated a large-scale paired multi-omics dataset encompassing whole-exome sequencing (488 samples), RNA-seq (470 samples), whole-genome bisulfite sequencing (363 samples), and microRNA sequencing (332 samples). Using these paired multi-omics data, we identified six spatial subtypes defined by the composition of 11 spatially annotated cellular compartments, reflecting distinct molecular profiles and immune infiltration states. In CC, HPV infection was associated with suppression of TGFβ signaling, disrupting spatial fibroblast–epithelial proximity and fostering immune-infiltrated TMEs. Across OC, EC, and CC, we delineated five malignant cell states characterized by distinct CNV patterns and oncogenic pathway activation, contributing to immune evasion. Focusing on HGSOC, we identified a proliferative malignant niche enriched for CIN-high cells, marked by enhanced proliferation and oxidative metabolism, immune exclusion, and strong associations with recurrence. This niche engaged in extensive crosstalk with fibroblasts and macrophages, jointly promoting immune evasion and chemoresistance through stromal protection and immune suppression. Leveraging these insights, we developed a predictive framework based on a pathology foundation model applied to hematoxylin and eosin (H&E)-stained slides to estimate niche abundance. Application of this model to two independent cohorts enabled accurate prediction of patient survival and bevacizumab response, demonstrating both the prognostic and translational utility of spatially defined tumor niches. To facilitate broader access, the processed dataset is publicly available via STAGE (http://stage.pku-genomics.org/), a user-friendly platform for visualization and data access.

## Results

### Cohort design and multi-omics data generation for FGTs

To systematically investigate genetic alterations, molecular dysregulation, and their TME interactions in FGTs, we assembled a large-scale cohort comprising 546 tumor samples from 477 patients, collected before or after neoadjuvant therapy (**Fig. 1a**). The cohort was annotated with detailed clinical metadata, including physical characteristics, diagnosis, treatment history, therapeutic response, and prognosis (**Supplementary Table 1**), with summary statistics provided across cancer types (**Supplementary Table 2**). Specifically, the cohort included 290 samples from 229 ovarian cancer (OC) patients, 140 samples from 136 endometrial cancer (EC) patients, 94 samples from 94 cervical cancer (CC) patients, and 22 samples from 18 patients with other gynecologic tumors (**Fig. 1b**). More than 80% of OC cases were diagnosed at advanced stage (III or IV), and nearly half experienced relapse (**Supplementary Table 2**). In contrast, recurrence was less frequent in EC (33%) and CC (19%).

**Fig. 1:**
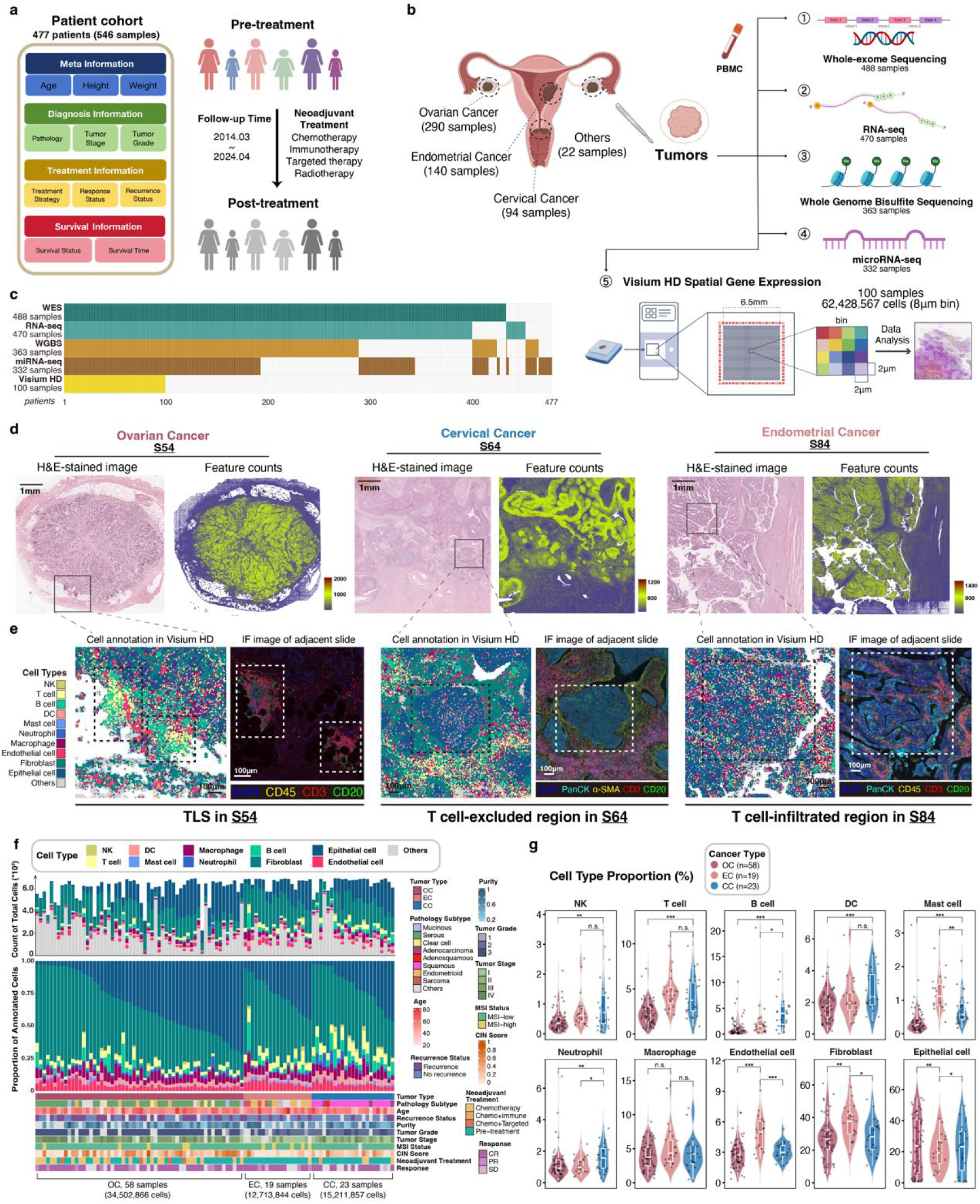
Cohort design and spatial transcriptomics profiling of female genital tumors using Visium HD with multi-omics integration. **a,** Schematic of patient enrollment and treatment in the female genital tumor (FGT) cohort. **b,** Collection of tumor and blood samples for multi-omics profiling, including paired whole-exome sequencing (WES), RNA sequencing (RNA-Seq), whole genome bisulfite sequencing (WGBS), microRNA sequencing (miRNA-seq), and Visium HD spatial transcriptomics. **c,** Overview of patients whose paired samples successfully sequenced across multiple platforms. **d,** Spatial maps of three representative tumors (different cancer types), analyzed with 8 µm bins. Shown are H&E-stained images (left) and spatial feature counts (right). **e,** Magnified regions of interest (ROIs) from (**d)** showing cell-type annotations from Visium HD data alongside matched immunofluorescence (IF) images (adjacent slides). **f,** Sample-wise summary of total cell counts (top), cell-type composition (middle), and clinical information (bottom) in the spatial transcriptomics dataset. **g,** Distribution of cell-type proportions across cancer types. Wilcoxon rank-sum test, ***p<0.001, **p<0.01, *p<0.05.

Multi-omics profiling was performed on collected tumor samples, including whole-exome sequencing (WES, for matched blood samples as well), RNA sequencing (RNA-seq), whole-genome bisulfite sequencing (WGBS), microRNA sequencing (miRNA-seq), and spatial transcriptomics (**Fig. 1b,c**). To address the resolution limitations of conventional spatial transcriptomics platforms, we utilized the 10x Visium HD platform to profile 100 formalin-fixed paraffin-embedded (FFPE) tumor sections from 99 patients, integrated with the other four omics datasets. Detailed patient-sample pairing information is provided (**Supplementary Table 3**).

### High-resolution spatial transcriptomics reveals cellular composition, microstructural features, and immune infiltration in FGTs

To characterize spatial heterogeneity across FGTs, we generated high-resolution spatial transcriptomics data from 100 FFPE sections using Visium HD, achieving single-cell resolution (2 μm binning) and processed at 8 μm for downstream analysis. In total, 62,428,567 cells were obtained across the 100 sections. The Visium HD dataset yielded an average of 800-1,000 detected genes per spot within tumor regions, indicating high-quality sequencing data (**Fig. 1d**).

We developed a robust cell-type annotation pipeline that integrated spatial expression profiles with marker-gene signatures to assign cell-type scores (**Supplementary Table 4**). This approach enabled the accurate identification of distinct microenvironmental structures, including tertiary lymphoid structures (TLS), T cell-excluded tumor regions, and T cell-infiltrated tumor regions, validated by paired H&E staining and immunofluorescence (IF) from adjacent sections (**Fig. 1e and Extended Data Fig. 1a-c**). Spatial localization of major cell types corresponded with expected marker expression patterns, *CD3E*, *CD3D*, *TRAC*, *GNLY*, *MS4A1*, *CD14*, *CD68*, and *ITGAX* for immune cells, *COL1A1* for fibroblasts, and *PECAM1* for endothelial cells (**Extended Data Fig. 1c-g**). Marker-specific expression patterns were consistent across all 100 samples, supporting annotation reliability (**Extended Data Fig. 1h**).

Quality-control analyses showed that epithelial cells exhibited the highest overall transcript counts, including mitochondrial transcripts (**Extended Data Fig. 1i**). To validate annotation accuracy further, we performed cosine similarity analysis on 80,000 randomly selected cells (800 per sample), which demonstrated strong within–cell type similarity (**Extended Data Fig. 1j**). To validate the sequencing accuracy of Visium HD, we compared the ST expression profiles with paired RNA-seq data of the 100 tumors. pseudo-bulk expression profiles were generated by summing all spatial bins per sample and normalized to transcripts per million (TPM). Pearson correlation coefficients (PCCs) between pseudo-bulk Visium HD profiles and matched RNA-seq revealed high concordance, with median PCC values >0.60 across 100 samples (**Extended Data Fig. 1k**). Immune infiltration estimates derived from RNA-seq deconvolution using CIBERSORT^21^ were also significantly correlated with immune cell proportions annotated from Visium HD (**Extended Data Fig. 1l**). Gene-level comparisons demonstrated high cross-platform concordance across 16,926 shared genes (**Extended Data Fig. 1m**).

Analysis of cell-type proportions across the 100 sections revealed heterogeneous TMEs (**Fig. 1f**), with fibroblasts and epithelial cells as the predominant populations (**Extended Data Fig. 1n**). Notably, OC samples exhibited the lowest immune infiltration and highest epithelial fraction (**Fig. 1f,g**), consistent with RNA-seq–based deconvolution (**Extended Data Fig. 1o**) and previous reports describing OC as an “immune-cold” tumor type^19^.

### Identification of spatial cellular subtypes and their molecular clusters in FGTs

To resolve molecular heterogeneity across FGTs, we first performed unsupervised non-negative matrix factorization (NMF) consensus clustering of RNA-seq data from 250 OC, 124 EC, and 84 CC samples. This analysis identified six distinct molecular clusters (**Extended Data Fig. 2a**). Marker gene expression revealed clear differences in immune, epithelial, and stromal signatures, delineating immunophenotypic profiles for each cluster. Among serous subtypes, samples segregated into two clusters with contrasting immune and proliferative features, termed *Low-proliferative serous-enriched* and *High-proliferative serous-enriched*, corresponding to the previously described *Immunoreactive* and *Proliferative* clusters in HGSOC^12,22^. A separate *Stromal-rich* cluster was defined by high stromal marker expression. Non-serous subtypes, particularly endometrioid tumors from OC and EC, were grouped into a *Serous-depleted* cluster, underscoring their transcriptional divergence from serous tumors. Most CC samples were classified into a *CC-enriched* cluster with elevated immune and proliferation-associated gene expression, reflecting high immune infiltration in CC.

To map spatial architecture, we next applied unsupervised clustering to 62,428,567 cells profiled by Visium HD across 100 FGT samples (**Fig. 2a**). This analysis identified 11 distinct cellular compartments. which were classified based on enrichment of epithelial, immune, and stromal cell types (**Extended Data Fig. 2b**). Using compartment proportions per sample, hierarchical clustering defined six spatial cellular subtypes: (1) Epithelium-enriched, (2) Immune-restricted, (3) Necrotic, (4) Angio-fibrotic, (5) Immune-stromal, and (6) Lymphoid-enriched (**Fig. 2b and Extended Data Fig. 2b**). Subtype distribution differed across cancer types: CC samples were enriched in Immune-stromal and Lymphoid-enriched subtypes, whereas OC samples clustered predominantly in Epithelium-enriched and Immune-restricted subtypes, reflecting “immune-hot” versus “immune-cold” phenotypes, respectively.

**Fig. 2:**
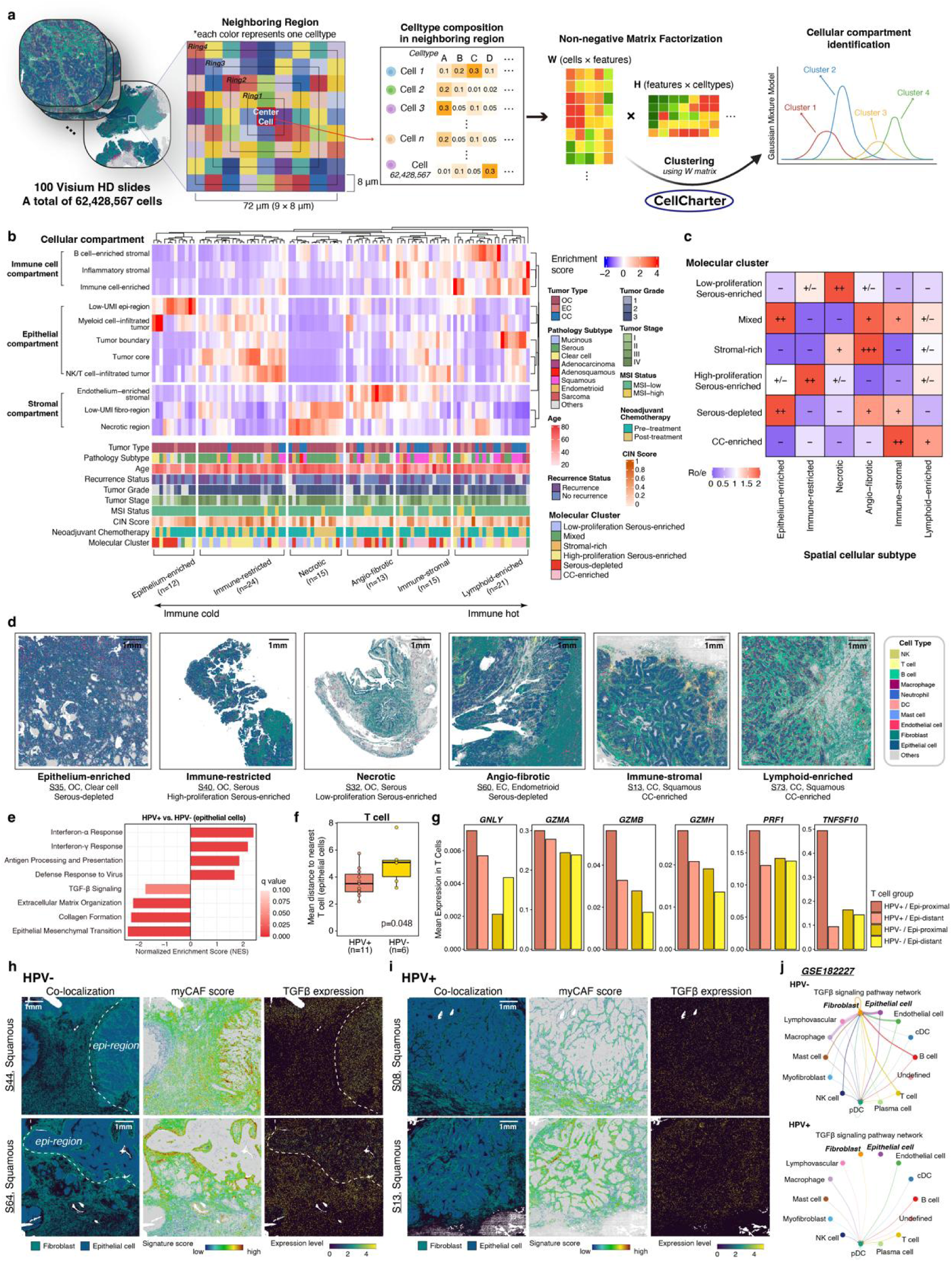
Spatial cellular subtypes, molecular features, and immunophenotypes in the FGT cohort. **a,** Workflow for identifying spatial cellular compartments. **b,** Hierarchical clustering of cellular compartment composition across 100 samples with clinical annotation, yielding six distinct spatial cellular subtypes with increasing immune infiltration. **c,** Molecular enrichment of each spatial subtype, summarized as the ratio of observed to expected sample frequency (Ro/e). +++, Ro/e > 3; ++, 2 ≤ Ro/e ≤ 3; +, 1.2 ≤ Ro/e < 2; +/−, 0.8 ≤ Ro/e < 1.2; −, Ro/e < 0.8. **d,** Representative samples from each spatial subtype. **e,** GSEA comparing epithelial cells from HPV-positive versus HPV-negative CC samples. **f,** Mean epithelial-to-nearest-T-cell distance across slides stratified by HPV-infection status in CC samples. Significance was assessed by the Wilcoxon rank-sum test. **g,** Mean expression level of cytotoxicity-related genes across T-cell groups, stratified by HPV status and distance to epithelial cells in CC samples. **h,i,** Spatial distribution of fibro-epithelial co-localization, myCAF signature score, and TGFβ expression (*TGFB1*+*TGFB2*+*TGFB3*) in representative HPV-negative (**h**) and HPV-positive (**i**) CC samples. **j,** HPV-dependent modulation of TGFβ signaling interactions across cell types in public scRNA-seq datasets analyzed with CellChat.

Integration of spatial and molecular classifications revealed distinct molecular feature enrichments within each spatial subtype (**Fig. 2c**). The Immune-restricted subtype correlated with the High-proliferative serous-enriched cluster, consistent with reduced immune gene expression, while the Necrotic subtype aligned with the Low-proliferative serous-enriched cluster, characterized by downregulated proliferation. As expected, CC-enriched molecular clusters corresponded to Immune-stromal and Lymphoid-enriched subtypes, confirming enhanced immune infiltration in CC. Collectively, these analyses demonstrated that molecular clusters correspond to specific spatial architectures, linking transcriptional programs to defined microenvironmental patterns. For instance, tumors with high immune scores (e.g., CC-enriched samples S13 and S73) exhibited spatial co-localization of T cells with epithelial cells, in contrast to tumors with low immune scores (e.g., serous-enriched samples S35 and S40) (**Fig. 2d**).

### HPV infection mediates inflammatory responses in CC through TGF-β suppression

Based on spatial subtype observations, we aim to determine factors contributing to elevated T cell infiltration in CC samples. High-risk human papillomavirus (HPV) infection is a major driver of CC and has been linked to increased tumor-infiltrating lymphocytes^23^. In our cohort, genome-wide analysis of 80 CC samples identified 10 single-base substitution (SBS) mutational signatures (**Extended Data Fig. 2c and Supplementary Table 5**), seven of which matched the COSMIC SBS signatures^24^. Three novel signatures (SBS-CC1, SBS-CC2, and SBS-CC3) were identified in 73, 12, and 47 tumors, respectively (**Extended Data Fig. 2c**). SBS2 and SBS13, associated with APOBEC mutagenesis^24^, frequently co-occurred in CC tumors (**Extended Data Fig. 2d**). In parallel, WGBS revealed hypomethylation of the *APOBEC3C* promoter in CC relative to OC and EC (**Extended Data Fig. 2e**), while Visium HD confirmed upregulation of APOBEC3 family genes in CC (**Extended Data Fig. 2f**), collectively highlighting HPV-associated activation of APOBEC3 enzyme activity. HPV genotyping by qPCR classified 17 Visium HD CC samples as HPV-positive (n=11) or HPV-negative (n=6) (**Supplementary Table 6**), with APOBEC-associated signatures enriched in HPV-positive tumors (**Extended Data Fig. 2g**).

Pathway enrichment analysis demonstrated that CC epithelial cells exhibited activation of cell cycle and immune pathways (**Extended Data Fig. 3a**), consistent with HPV-driven transcriptional programs^25^. In particular, HPV-positive epithelial cells displayed elevated type I interferon (IFN) and antigen presentation signaling (**Fig. 2e**), reflecting antiviral immune activation. Viral antigens such as HPV16/18 E6/E7 proteins are recognized by CD8+ T cells^26^, supporting our observation that T cells were positioned closer to epithelial cells in HPV-positive compared to HPV-negative tumors (**Fig. 2f**). Although T-cell abundance did not differ significantly between groups (**Extended Data Fig. 3b**), spatially proximal T cells in HPV-positive tumors expressed higher levels of cytotoxic effector genes (**Fig. 2g**). Consistently, APOBEC-associated CC samples showed higher IFN-γ signaling (IFNG) and cytotoxic T lymphocyte (CTL) signature scores (**Extended Data Fig. 3c**).

Spatial analysis further revealed that HPV-positive tumors exhibited epithelial-T cell co-localization with CXCL10 upregulation (**Extended Data Fig. 3d,e**), a chemokine that recruits effector lymphocytes and promotes “hot” TMEs^27^. Strikingly, epithelial cells in HPV-positive tumors showed reduced TGFβ and EMT pathway activity (**Fig. 2e and Extended Data Fig. 3e**), consistent with prior studies in HPV-driven head and neck cancers^28,29^. In HPV-negative tumors, myofibroblasts (myCAFs) surrounded epithelial regions, supported by elevated TGFβ signaling (**Fig. 2h**), forming barriers that excluded T cells (**Extended Data Fig. 3f**). In contrast, HPV-positive tumors displayed reduced myCAF scores and TGFβ signaling, allowing T cells to infiltrate epithelial regions and exert cytotoxic activity (**Fig. 2i, Extended Data Fig. 3g**). IF images validated these findings: α-SMA+ myCAF accumulation was evident in HPV-negative sample S64, whereas HPV-positive sample S13 exhibited reduced myCAF enrichment and dense immune infiltration (**Extended Data Fig. 3h,i**).

To generalize these findings, we analyzed public scRNA-seq data from HPV-positive and HPV-negative oropharyngeal squamous cell carcinomas^25^. Cell–cell interaction analysis showed elevated epithelial–T cell communication via MHC-I ligand–receptor pairs in HPV-positive tumors, while epithelial–fibroblast interactions mediated by TGFβ were reduced (**Fig. 2j and Extended Data Fig. 3j,k**). Together, these results demonstrate that HPV infection suppresses TGFβ signaling, disrupts myCAF recruitment, and thereby enhances T cell infiltration and cytotoxicity in CC.

### CIN heterogeneity and immune phenotypes across pathology subtypes

Building on the distinct molecular and spatial architectures observed between serous and non-serous tumors (**Fig. 2b,c, Extended Data Fig. 2a**), we next examined the mechanisms underlying subtype-specific genomic and microenvironmental features in OC and EC, focusing on serous and endometrioid subtypes.

Whole-exome sequencing revealed divergent genomic landscapes. Serous tumors harbored frequent TP53 mutations and widespread chromosomal gains and losses, whereas endometrioid tumors exhibited higher rates of microsatellite instability (MSI-high), lower CIN levels, and recurrent mutations in *ARID1A*, *PTEN*, *CTNNB1*, and *PIK3CA* (**Fig. 3a,b**). These findings are consistent with prior studies reporting high CIN in serous but not endometrioid tumors^9^, a pattern corroborated by our cohort (**Extended Data Fig. 4a**). In line with previous reports linking CNV burden to reduced immune infiltration^30^, CIN scores were negatively correlated with T-cell scores in both serous and endometrioid tumors based on paired WES and RNA-seq data (**Extended Data Fig. 4b**). Spatial transcriptomics further revealed higher proportions of NK and T cells in endometrioid tumors, whereas epithelial cell proportions were comparable across subtypes (**Fig. 3c**), highlighting increased immune infiltration in endometrioid tumors.

**Fig. 3:**
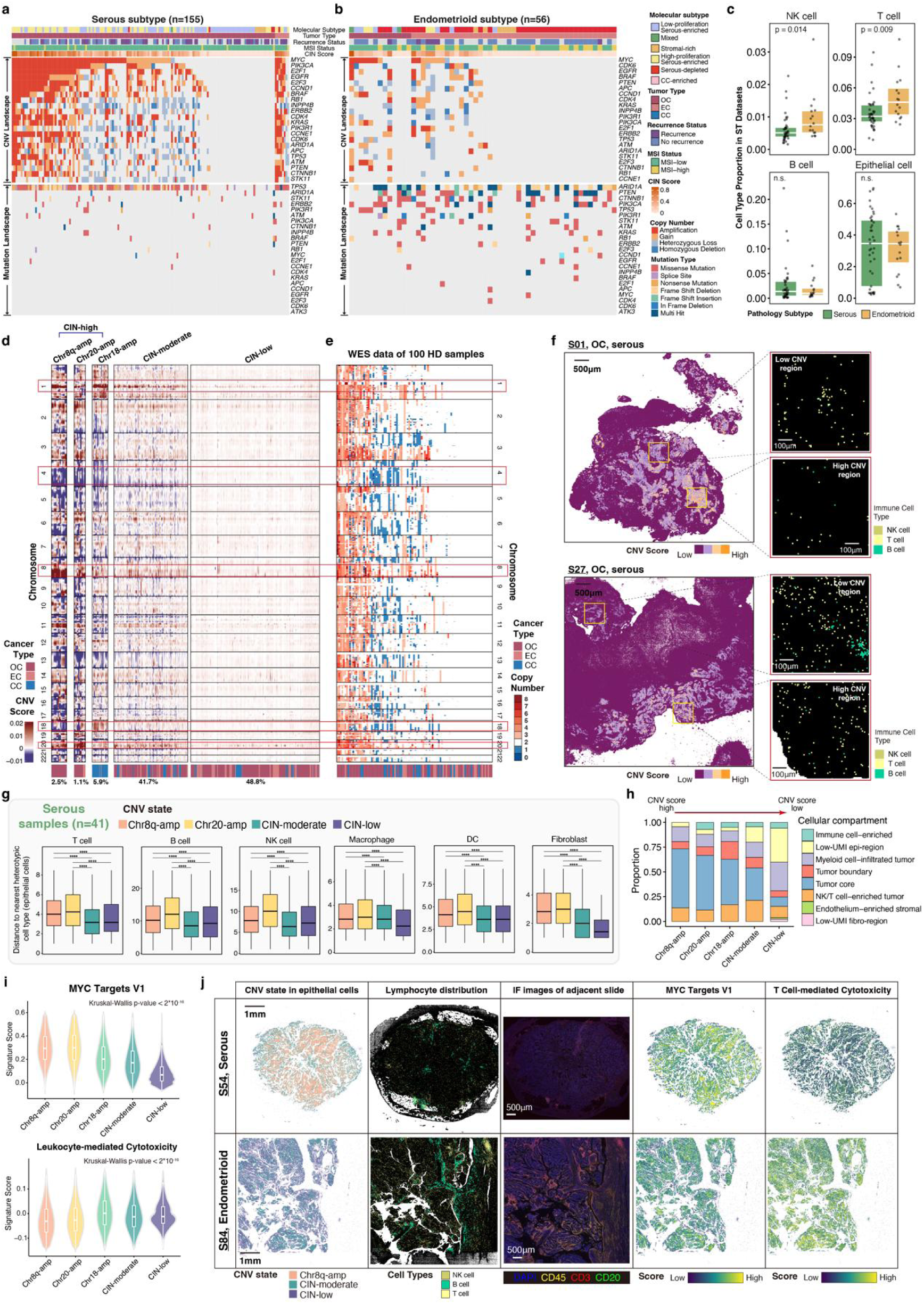
Spatial CNV patterns and tumor heterogeneity in the FGT cohort. **a,b,** OncoPrints showing CNV and mutation profiles of key driver genes in serous subtype (**a**) and endometrioid subtype (**b**) samples. **c,** Comparison of cell-type proportions between serous and endometrioid subtypes in the Visium HD dataset. Each dot represents one sample. Wilcoxon rank-sum test. **d,** Clustering of spatial CNV profiles across 100 FGT samples, inferred from Visium HD spatial transcriptomics data. CIN, chromosomal instability. **e,** CNV patterns derived from matched WES for the same samples shown in (**d**). **f,** Representative ROIs showing spatial variation of CNV scores (left) alongside lymphocyte infiltration levels (right). **g,** Distance between epithelial cells and nearest different cell type in serous tumors, stratified by CNV state. Wilcoxon rank-sum test, ****p<0.0001. **h,** Relative proportion of epithelial cells within cellular compartments across five CNV states. **i,** Expression of cancer-related pathways stratified by CNV state. P values from Kruskal-Wallis test. **j,** Representative spatial maps of CNV states, lymphocytes infiltration, IF images of adjacent slide, MYC pathway activity, and T cell cytotoxicity in serous (top) and endometrioid (bottom) tumors.

To dissect intratumoral CNV heterogeneity at high resolution, we applied inferCNV^31^ to 100 Visium HD samples, generating genome-wide CNV profiles. As expected, epithelial cells exhibited the highest CNV burden (**Extended Data Fig. 4c,d**), with multiple CNV-defined clusters per tumor section (**Extended Data Fig. 4d-f**), reflecting spatially distinct tumor subclones. Across all samples, epithelial cells were classified into three major categories by CIN extent: CIN-high, CIN-moderate, and CIN-low (**Fig. 3d**). Within CIN-high tumors, we identified three recurrent focal amplification states, including Chr8q-amp, Chr20-amp, and Chr18-amp. We used *MYC* (8q24.21) amplification status as a metric for CIN evaluation. Consistent with our findings, inferred CIN-high regions exhibited higher gene amplification prevalence after MYC probe optimization (**Extended Data Fig. 4g**), while cells within these regions showed increased *MYC* amplification frequency (**Extended Data Fig. 4h**). Validation against paired WES confirmed concordant CNV profiles, including recurrent alterations on chromosomes 1q, 4, 8, 14, and 20 (**Fig. 3d,e**), with high similarity between Visium HD– and WES-derived profiles (**Extended Data Fig. 4i**). CNV scores decreased stepwise from CIN-high to CIN-low, and were near zero in normal cells (**Extended Data Fig. 4j**). Subtype-specific enrichment was observed: Chr8q-amp and Chr20-amp were frequent in serous tumors, while Chr18-amp was predominant in CC (**Fig. 4d and Extended Data Fig. 4k**). In contrast, endometrioid tumors showed minimal CIN-high states, consistent with their lower CNV burden (**Extended Data Fig. 4k and Supplementary Table 7**), consistent with the reduced CNV frequencies in the endometrioid cohort (**Fig. 3a,b**).

**Fig. 4:**
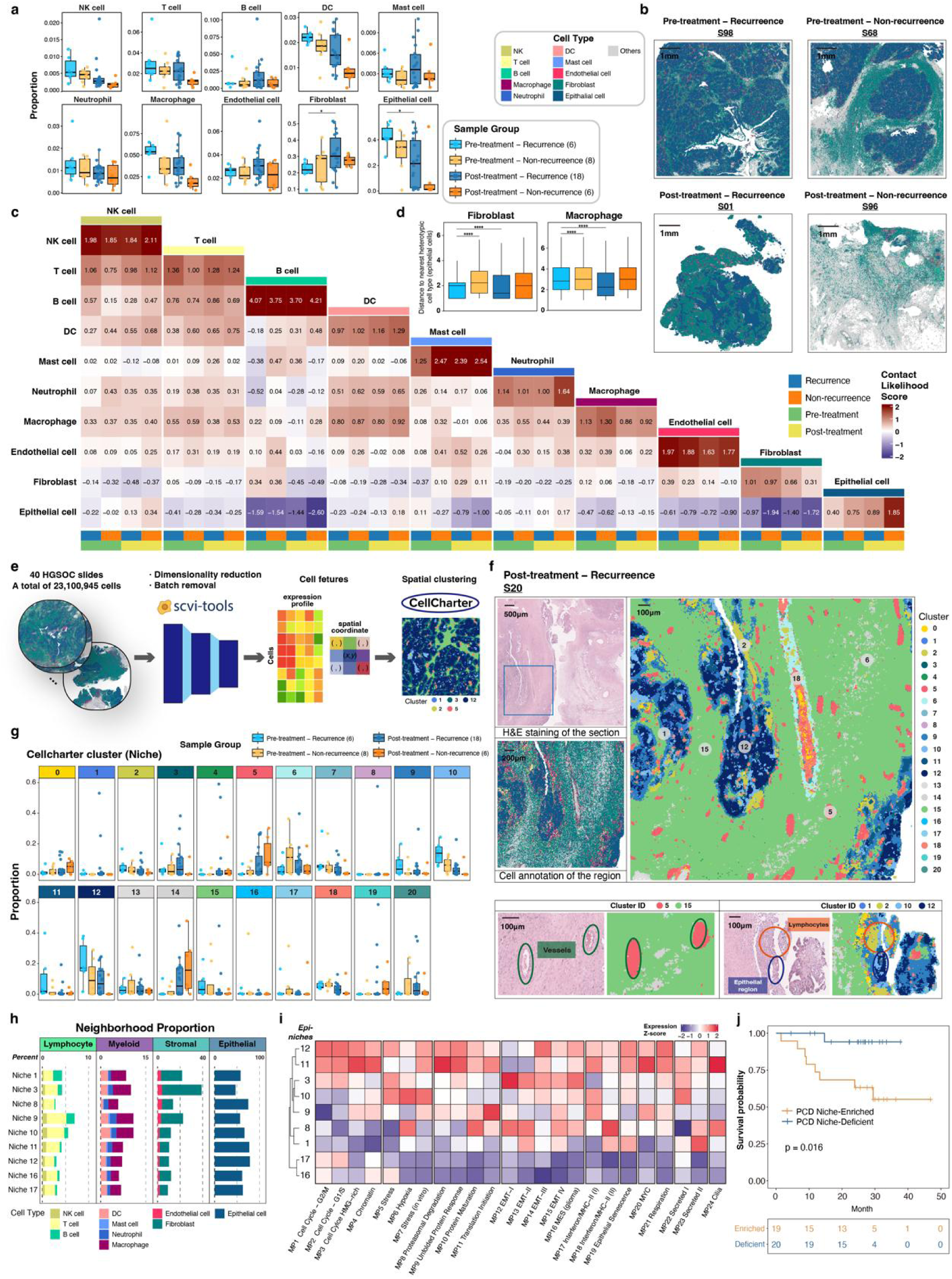
Spatial organization and molecular features of tumor niches in high-grade serous ovarian cancer (HGSOC). **a,** Cell-type distributions in HGSOC samples, stratified by treatment and recurrence status. Each dot represents one sample. Wilcoxon rank-sum test, *0.01≤p<0.05. **b,** Representative spatial maps of annotated cells in samples (S98, S68, S01, and S96) from different treatment and recurrence groups. **c,** Heatmap of predicted cell-cell interaction likelihood across cell types by treatment and recurrence groups. **d,** Mean epithelial-to-nearest fibroblasts (left) or macrophage (right) distance across treatment and recurrence groups. Wilcoxon rank-sum test, ****p<0.0001. **e,** Workflow for spatial clustering across 40 HGSOC samples. **f,** Representative sample of post-treatment recurrent sample (S20), showing H&E image (top left), spatial annotation (middle left), and spatial clustering (right). Spatial clustering of ROIs is displayed with H&E morphology (bottom). **g,** Distribution of annotated spatial clusters across treatment and recurrence groups. **h,** Proportions of cell types surrounding epithelial cells across epithelial niches. **i,** Expression of 24 recurrent malignant metaprograms (MPs) across epithelial niches. **j,** Kaplan-Meier survival analysis stratified by median abundance of the Proliferating Core Desert (PCD) niche. Log-rank test.

Previous analysis linked CNV states with immune exclusion (**Extended Data Fig. 4b**). In representative HGSOC samples (S01, S27), lymphocyte infiltration was markedly reduced in high-CNV regions compared to low-CNV regions (**Fig. 3f**). IF image confirmed absent T cell distribution in high CNV region, whereas elevated T cell infiltration is observed in low CNV region within the same tissue slide (**Extended Data Fig. 4l**). Quantitative analysis revealed that CIN-high epithelial cells were located further from immune and stromal cells than CIN-moderate or CIN-low states (**Fig. 3g**). Similar patterns were observed in endometrioid and CC tumors for Chr8q-amp and Chr20-amp, although they were less frequent in non-serous tumor (**Extended Data Fig. 4m,n**). By contrast, Chr18-amp epithelial cells showed distinct co-localization with immune cells, differing from other CIN-high states. At the population level, CIN-high epithelial cells localized predominantly to tumor cores, while CIN-moderate and CIN-low states were enriched in immune-infiltrated regions (**Fig. 3h**). These findings support a spatially resolved negative relationship between CNV burden and immune infiltration, consistent with the “immune-cold” phenotype of HGSOC. By contrast, EC and CC samples, which had lower CIN scores, displayed enhanced lymphocyte infiltration (**Extended Data Fig. 4o**).

### CNV-driven oncogenic pathways and immune evasion in serous tumors

To explore functional consequences of CNVs, we examined oncogene (OG) and tumor suppressor gene (TSG) scores^32^. Mean CNV scores correlated strongly with OG probabilities (p<0.001) (**Extended Data Fig. 5a**), suggesting selection pressure on functionally relevant alterations. CIN-high states were enriched for amplifications in *JUN* (1p32.1), *PIK3CA* (3q26.32), *MYC* (8q24.21), *E2F1* (20q11.22), and *SRC* (20q11.23), and deletions in *FAT1* (4q35.2) (**Extended Data Fig. 5b**). These findings were validated through WES datasets, as *MYC* and *PIK3CA* amplifications were frequent in serous tumors (>60% and >50%, respectively) but rare in endometrioid (<25%) (**Fig. 3a,b**). Overall, ST datasets revealed a strong positive correlation between *MYC* expression levels and *MYC* CNV scores across serous and endometrioid subtypes (**Extended Data Fig. 5c**), with *MYC* showing strong amplification and expression within epithelial compartments spatially (**Extended Data Fig. 5d**). By contrast, *CDK4* showed limited correlation between gene amplification and expression (**Fig. 3a,b and Extended Data Fig. 5d**).

As prior studies reported that elevated MYC signaling and PI3K signaling pathways mediate T cell exclusion in tumors^33,34^, pathway analysis revealed that CIN-high states, particularly Chr8q-amp and Chr20-amp, exhibited the highest activation of MYC and PI3K signaling, coupled with the lowest immune pathway activity (**Fig. 3i and Extended Data Fig. 5e**). Spatially, we identified a unique spatial pattern of lymphocytes, which were localized exclusively outside CIN-high regions (**Fig. 3j**). These excluded regions demonstrated elevated MYC signaling activity and were predominantly observed in serous tumors, with reduced immune-mediated tumor cell killing due to their immune-excluded phenotypes (**Fig. 3j**). Differentially methylated region analysis between serous and endometrioid subtypes revealed that *PIK3R1*, a negative regulator of the PI3K signaling pathway, was aberrantly silenced in serous tumors, resulting in reduced gene expression levels (**Extended Data Fig. 5f**), while miRNA profiling revealed dysregulation of several miRNAs affecting MYC and PI3K pathways, including miR-375-3p (tumor suppressive)^35^ and miR-1269b (oncogenic)^36^ (**Extended Data Fig. 5g**). Together, these results demonstrate that CIN-high subclones in serous tumors drive immune exclusion through amplification and activation of oncogenic programs, particularly MYC and PI3K signaling, thereby promoting tumor progression and immune evasion.

### Characterization of spatial cellular interactions and intratumor heterogeneity in HGSOC

Given the immune-excluded phenotypes and spatial CIN heterogeneity observed in serous tumors, we systematically analyzed intratumoral transcriptomic heterogeneity and cell–cell interactions in 40 HGSOC samples profiled by Visium HD. Patients were stratified into four clinical subgroups by neoadjuvant chemotherapy and recurrence status. Chemotherapy markedly reduced epithelial cell proportions, particularly in non-recurrence groups, where epithelial regions were nearly eliminated, suggesting effective tumor clearance (**Fig. 4a and Extended Data Fig. 6a**). In contrast, recurrence groups retained epithelial-rich regions, while post-treatment non-recurrence samples displayed extensive necrotic areas classified as “Others” cell types (**Fig. 4b**).

We next quantified spatial interactions using cell–cell neighboring scores. Strong intra-immune cell interactions were observed, while epithelial cells were generally spatially excluded from T and B cells (**Fig. 4c**). Chemotherapy partially enhanced immune–epithelial co-localization, particularly with macrophages, consistent with previous reports^34^. Distance analyses revealed that fibroblasts and macrophages were significantly closer to epithelial cells in recurrence groups, with the shortest distances in the post-treatment recurrence subgroup (**Fig. 4d**). Macrophage infiltration into epithelial regions was particularly pronounced in recurrence samples (**Extended Data Fig. 6b**), a pattern not observed for other immune populations (**Extended Data Fig. 6c**).

To further resolve tissue organization, we applied CellCharter^37^ to segment 40 HGSOC sections into spatial niches based on gene expression and local context (**Fig. 4e**). This analysis identified 21 clusters across all samples (**Extended Data Fig. 6d)**. Lymphocytes were enriched in niches 0 and 2, myeloid cells in niche 2, and endothelial–fibroblast aggregates in niches 5 and 18. Nine niches contained more than 30% epithelial cells (niches 1, 3, 8–12, 16, and 17), accounting for more than 95% of epithelial cells (**Extended Data Fig. 6e**). These epithelial niches varied in location: niche 3 was enriched for tumor boundaries with myeloid co-localization, niches 11 and 12 localized to tumor cores, and niches 16 and 17 contained low-UMI epithelial regions (**Extended Data Fig. 6f**). Differential gene expression confirmed niche-specific molecular profiles (**Extended Data Fig. 6g and Supplementary Table 8**), supporting the specific spatial organization of each niche. In post-treatment recurrence sample S20, more than 10 distinct niches aligned with histological domains such as vasculature, immune aggregates, and epithelial regions (**Fig. 4f**), supporting these niches represent conserved tissue domains across the HGSOC samples (**Extended Data Fig. 6h**). Niche frequencies differed by clinical status: the epithelial-rich niche 12 was more abundant in recurrence groups, whereas necrotic niche 14 was enriched post-treatment (**Fig. 4g**).

### The Proliferating Core Desert (PCD) niche: a malignant program linked to immune exclusion

We next focused on epithelial niches to identify malignant programs associated with recurrence. Spatial neighborhood analysis revealed fibroblast enrichment around epithelial cells in niche 3, consistent with its boundary localization, while niche 12 contained predominantly epithelial neighbors and minimal lymphocyte infiltration, in line with its tumor-core location (**Fig. 4h**, **Extended Data Fig. 6f**). We next quantified the expression levels of previously defined gene-expression programs in cancer cells (meta-programs, MPs)^38^ across these epithelial niches (**Fig. 4i**). We observed that epithelial cells within Niche 12 exhibited elevated activity in cell cycle pathways, as well as MPs associated with MYC signaling and respiration process, suggesting enhanced proliferative capacity in this niche. Furthermore, CIN-high states were also concentrated within this niche (**Extended Data Fig. 6i-l**). Given its low immune infiltration, proliferative programs, tumor-core localization, and CIN-high enrichment, we designated niche 12 as the Proliferating Core Desert (PCD) niche. The PCD niche was more frequent in recurrence groups (**Fig. 4g**) and significantly larger in recurrent versus non-recurrent tumors (**Extended Data Fig. 7a**).

Differential expression analysis revealed that PCD niche epithelial cells upregulated immune-evasion and stromal-interaction genes, including *CD24* and *CD47* (encoding anti-phagocytic “don’t eat me” signals)^39,40^, as well as *WNT7A* and *MDK* (implicated in CAF-mediated aggressiveness)^41,42^ (**Extended Data Fig. 7b and Supplementary Table 9**). Gene set enrichment analysis confirmed activation of proliferation, MYC signaling, and PI3K signaling pathways, with downregulation of inflammatory signaling (**Extended Data Fig. 7b, c**). CIN-high enrichment, *MYC*/*PIK3CA* amplification, and elevated signaling scores were specifically observed in the PCD niche (**Extended Data Fig. 7d,e**), implicating CNV-driven oncogenic pathways in its immune-excluded phenotype.

Following chemotherapy, the PCD niche acquired additional resistance programs. Pathways related to EMT, glycolysis, oxidative metabolism, and WNT signaling were activated, while immune response signatures, including IFNα/γ and cytotoxicity, were attenuated (**Extended Data Fig. 7f**), which are hallmark features of chemoresistance cancer cells^43^. Thus, although chemotherapy increased immune infiltration overall (**Fig. 4c**), the PCD niche persisted as an immune-evasive domain, suggesting a role in tumor recurrence. Consistently, patients with higher proportions of PCD niche exhibited significantly reduced overall survival (**Fig. 4j**), establishing this niche as a prognostic determinant in HGSOC.

### Boundary CAFs act as protective neighbors of the PCD niche

To investigate how the PCD niche interacts with surrounding niches in HGSOC, we performed spatial co-localization analysis. The PCD niche was preferentially bordered by niche 3, enriched for CAFs at tumor boundary (**Fig. 4h**, **Fig. 5a, Extended Data Fig. 6f**). Spatial mapping further revealed that niche 3 often positioned between the PCD niche and fibroblast-rich niche 20 (**Fig. 5b**). Across 40 HGSOC samples, the PCD niche consistently co-occurred with niches 3 and 20 in the same sample (**Extended Data Fig. 8a**). Fibroblasts in niche 3 showed upregulation of extracellular matrix (ECM) remodeling pathways (**Extended Data Fig. 8b**), including collagens, matrix metalloproteinases (MMPs), fibronectin, syndecans, and SPP1 (**Extended Data Fig. 8c**), suggesting active TME remodeling to support tumor cell survival.

**Fig. 5:**
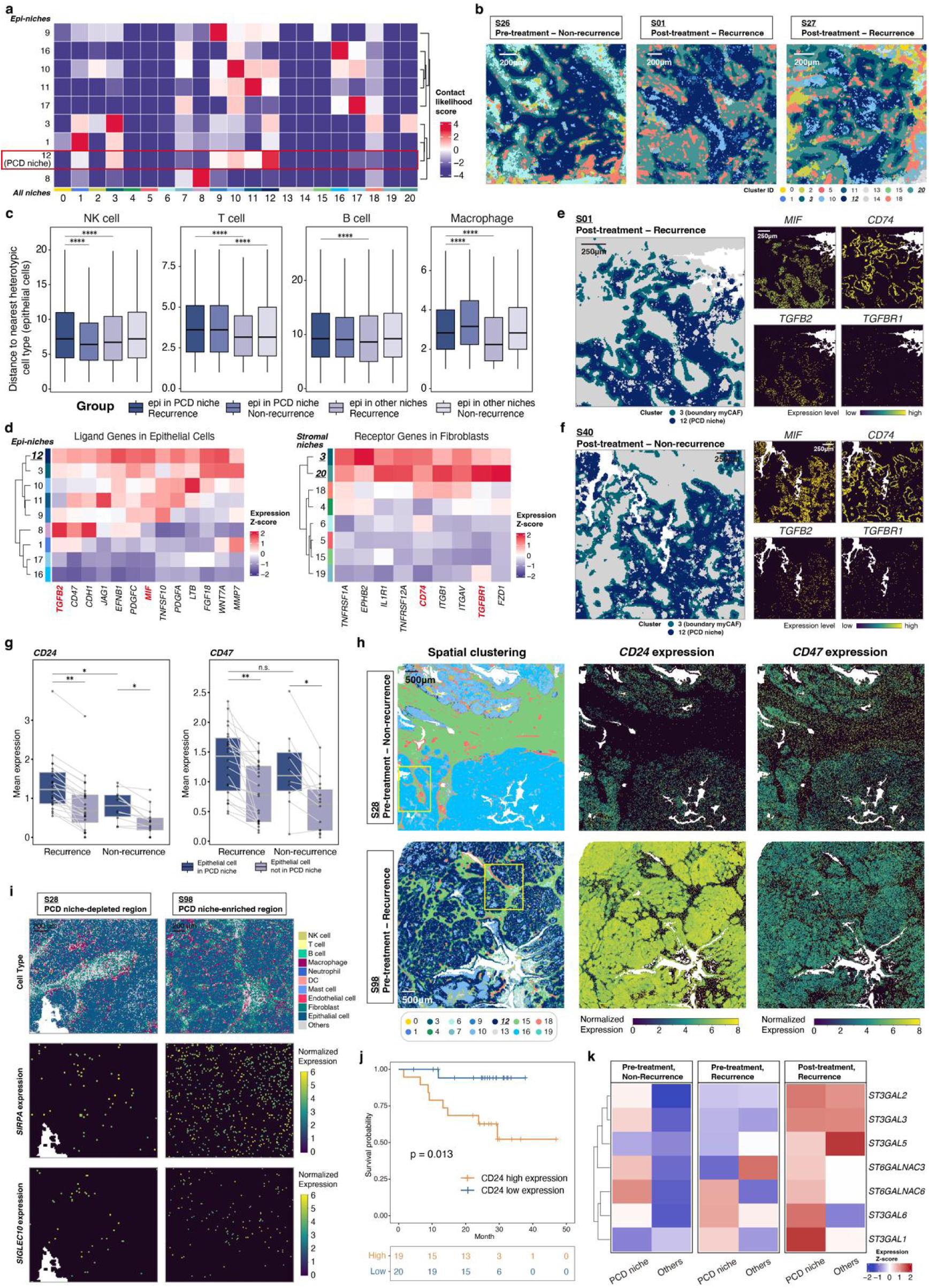
Immune evasion interactions between PCD niche and fibroblasts/macrophages in HGSOC. **a,** Heatmap of interaction likelihood between epithelial and other niches in HGSOC. **b,** Spatial co-localization of PCD niche with niches 3 and 20 across treatment and response groups. **c,** Distance between epithelial cells and nearest different cell type in HGSOC, stratified by epithelial niche location and recurrence status. Wilcoxon rank-sum test, ****p<0.0001. **d,** Heatmap of ligand expression in epithelial cells and receptor expression in fibroblasts, highlighting curated CAF-related ligand-receptor pairs. Visualization restricted to ligands upregulated in PCD (|log₂ FC| > 0.5, >1% bins) and corresponding receptors in niche 3 at the same criteria. **e,f,** Spatial mapping of PCD-niche 3 (left), interplay with ligand-receptor interactions of MIF-CD74 (top right) and TGFB2-TGFBR1 (bottom right) in S01 (**e**) and S40 (**f**). **g,** Mean *CD24* and *CD47* expression in epithelial cells inside versus outside PCD, stratified by recurrence status. Wilcoxon rank-sum test, **p<0.01, *0.01≤p<0.05, n.s. non-significant. **h,** Spatial maps of clusters (left) and normalized *CD24* (middle) and *CD47* (right) expression in non-recurrence (S28, top) and recurrence (S98, bottom) pre-treatment samples. **i,** Spatial characterization of cell type (top) and normalized expression of SIRPA (receptor of CD47, middle) and SIGLEC10 (receptor of CD24, bottom) in ROIs from recurrence (S98, left) and non-recurrence S28 (right) samples. **j,** Kaplan–Meier survival analysis stratified by median *CD24* expression. Log-rank test. **k,** Heatmap of sialic acid-modifying enzyme gene expression in epithelial cells inside versus outside PCD niche, stratifiedd by treatment and recurrence groups.

Immune exclusion was particularly evident where the PCD niche was bordered by niche 3 CAFs (**Extended Data Fig. 8d**). Fibroblasts in niche 3 localized adjacent to PCD epithelial cells (**Extended Data Fig. 8e**), creating barriers that increased separation between immune cells and epithelial regions in the PCD niche compared with other epithelial niches (**Fig. 5c**). This CAF-mediated CTL exclusion correlated with recurrence (**Extended Data Fig. 8f**). myCAF scores were elevated in niches 3 and 20 (**Extended Data Fig. 8g**,**h**), supporting their roles in establishing fibroblast-rich protective zones around the PCD niche. Accordingly, we designated niche 3 as the Boundary myCAF niche and niche 20 as the Adjacent myCAF niche.

To uncover mechanisms of PCD–CAF communication, we examined ligand–receptor interactions between PCD epithelial cells and boundary fibroblasts using integrated spatial datasets^44^. These analyses revealed two ligand-receptor interactions, MIF-CD74 and TGFB2-TGFBR1 (**Fig. 5d and Supplementary Table 10**). Both interactions localized at PCD–CAF interfaces (**Fig. 5e,f**), suggesting functional roles in promoting fibroblast activation and immune exclusion. Notably, TGFβ signaling is a well-established driver of myCAF induction and immune exclusion^45^, while MIF-CD74 signaling has been implicated in immune evasion^46,47^ and in supporting CAF survival in pancreatic cancer^48^. These findings highlight previously underexplored CAF–tumor interactions in HGSOC.

### Enhanced "don’t eat me" signals in the PCD niche

In recurrent tumors, the PCD niche displayed marked macrophage infiltration (**Fig. 5c and Extended Data Fig. 9a**), confirmed by IF staining of adjacent sections (**Extended Data Fig. 9b**). Differential expression analysis showed that epithelial cells in the PCD niche significantly upregulated *CD24* and *CD47* (**Extended Data Fig. 7b**). These encode ligands for macrophage receptors Siglec-10 (*SIGLEC10*) and SIRPα (*SIRPA*), respectively, which suppress phagocytosis. Both *CD24* and *CD47* were elevated in PCD epithelial cells, with higher *CD24* expression in recurrent compared to non-recurrent tumors (**Fig. 5g**). Spatial mapping illustrated strong *CD24* and *CD47* expression in recurrence sample S98, in contrast to minimal expression in non-recurrence sample S28 (**Fig. 5h**). Co-localization of CD24+/CD47+ epithelial cells with SIGLEC10+ and SIRPA+ macrophages was observed in PCD-enriched regions of S98 but absent in S28 (**Fig. 5i**). Importantly, patients with higher *CD24* expression had significantly shorter overall survival (log-rank p=0.013) (**Fig. 5j**).

Because Siglec-10 preferentially binds sialylated CD24, we further examined sialylation gene expression. Genes encoding enzymes involved in sialoglycan biosynthesis, which are previously linked to poor cancer survival^49,50^, were upregulated in PCD epithelial cells compared with other niches (**Fig. 5k**), suggesting that enhanced sialylation may strengthen CD24-Siglec-10 interactions and promote immune suppression.

### EREG+ macrophages infiltrate the PCD niche as an immunosuppressive subtype

To define TAM heterogeneity, we applied Spotiphy^51^ with TAM signatures from a single-cell atlas of HGSOC^52^ to deconvolute spatial data. Gene markers confirmed accurate TAM decomposition (**Extended Data Fig. 9c**). Three major TAM subsets were identified, C3+, EREG+, and C1QA+ TAMs, together comprising >90% of TAMs (**Extended Data Fig. 9d**). Among these, only EREG+ TAMs differed significantly between recurrent and non-recurrent tumors, being enriched in recurrence (**Extended Data Fig. 9e**). EREG+ TAMs preferentially infiltrated the PCD niche, while C1QA+ TAMs were excluded from the epithelial region (**Extended Data Fig. 9f**, **g**). C3+ and VCAN+ TAMs showed no spatial preference. Overall, EREG+ TAMs were enriched in epithelial compartments, whereas C1QA+ TAMs were enriched in immune compartments (**Extended Data Fig. 9h**). Transcriptomic profiling further highlighted their divergent roles: EREG+ TAMs downregulated antigen presentation, leukocyte activation, and immune-response pathways, while C1QA+ TAMs upregulated antigen presentation and immune-activation pathways (**Extended Data Fig. 9i**). Together, these findings suggest that EREG+ and C1QA+ TAMs represent distinct polarization states shaped by niche-specific localization. In particular, EREG+ TAM infiltration into the PCD niche, combined with CD24/CD47-mediated “don’t eat me” signaling, establishes an immunosuppressive microenvironment that promotes tumor persistence and recurrence in HGSOC.

### Identification of the PCD niches on H&E images using a pathology foundation model

Computational pathology has emerged as a powerful approach to integrate artificial intelligence (AI) into clinical workflows. Leveraging the compatibility of Visium HD assays with FFPE tissue and H&E staining, we aligned spatial transcriptomic maps of PCD niches with corresponding H&E slides. The distinct PCD distribution observed in Visium HD could also be discerned in matched H&E-stained regions (**Fig. 4f**), suggesting its potential as a pathology-derived biomarker for prognosis.

To operationalize this, we developed POWER (Pathology-based Ovarian cancer-specific Workflow for Evaluating Risk), a machine-learning framework to predict PCD niche proportions directly from H&E whole-slide images (WSIs). POWER integrates UNI^53^, a vision transformer–based pathology foundation model pre-trained on >100 million tissue patches from 100,426 WSIs, with a multilayer perceptron (MLP) trained using PCD niche proportions derived from 40 HGSOC samples (**Fig. 6a**). For each cropped H&E patch, UNI generates a 1,024-dimensional feature vector, which the MLP uses to predict PCD niche content.

**Fig. 6:**
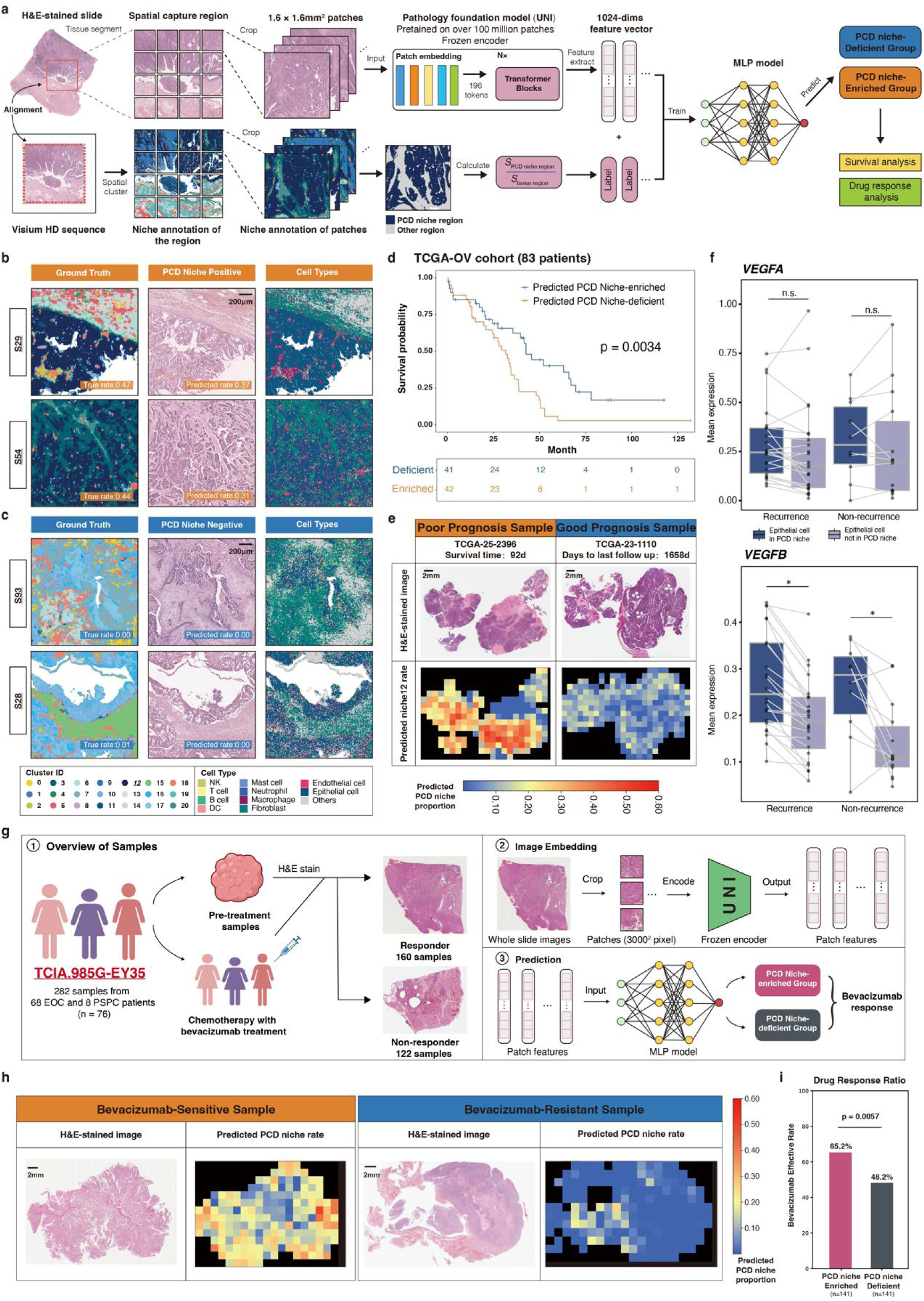
Prediction of PCD niche levels using pathology foundation models and clinical implications. **a,** Schematic workflow illustrating the multilayer perceptron (MLP) model predicting PCD niche levels from H&E-stained images, built on UNI, a foundation model pre-trained on >100,000 diagnostic whole-slide images (WSIs). Predictions were linked to prognosis (survival and therapy response). **b,c,** ROI-level predictions of PCD niche level compared with ground truth: high PCD (**b**) and low PCD (**c**). Panels show ground-truth PCD niche proportions (left), MLP predictions from H&E (middle), and spatial annotations (right). **d,** Kaplan-Meier survival analysis of the TCGA-OV cohort (n=83), stratified by predicted PCD niche levels from WSIs. Log-rank test. **e,** Representative TCGA-OV samples: poor prognosis (92 days, left) and good prognosis (1658 days, right). **f,** *VEGFA* and *VEGFB* expression in epithelial cells inside versus outside the PCD niche, stratified by recurrence. Wilcoxon rank-sum test. *0.01≤p<0.05, n.s. non-significant. **g,** Workflow showing the model to predict PCD niche levels and correlate with drug response in the TCIA.985G-EY35 cohort (76 OC patients treated with chemotherapy and bevacizumab). **h,** Representative WSI predictions from the TCIA.985G-EY35 cohort, showing a bevacizumab-sensitive (left) sample and a resistant (right) sample. **i,** Bevacizumab response rates stratified by predicted PCD niche levels (n=282, PCD-enriched vs. -deficient). Chi-Square test.

Model evaluation across multiple classifiers identified the MLP as the top performer (**Extended Data Fig. 10a**). With a 20% cutoff for PCD-positivity for classifying patches, POWER achieved an area under the curve (AUC) of 0.97, F1 score of 0.88, precision of 0.90, and recall of 0.86. Predictions correlated strongly with ground truth proportions in the test set (Pearson’s ρ = 0.90, p < 0.001; **Extended Data Fig. 10b**) and validation set (**Extended Data Fig. 10c,d**). POWER reliably distinguished PCD-positive from PCD-negative epithelial regions, despite both containing epithelial cells (**Fig. 6b,c**), confirming its specificity for identifying malignant epithelial states.

### POWER predicts prognosis and therapy response

To assess clinical utility, POWER was applied to the TCGA-OV cohort (n=83), where WSIs were paired with survival data. Patients stratified by predicted PCD niche abundance showed significantly worse survival in the high-PCD group (log-rank p=0.0034; **Fig. 6d**). Representative cases illustrated this association: a patient with poor prognosis (92 days survival) exhibited extensive PCD-positive regions, while a long-term survival patient (1,658 days) showed minimal PCD involvement (**Fig. 6e**). Non-tumor areas consistently had low predicted PCD scores, supporting model specificity (**Extended Data Fig. 10e**).

Molecular profiling revealed that epithelial cells within PCD niches exhibited hypoxia-related features, including elevated oxidative metabolism and respiration (**Fig. 4i, Extended Data Fig. 7b**). *VEGFB* expression was significantly higher in PCD epithelial cells, while *VEGFA* showed a non-significant trend (**Fig. 6f**). In sample S01 (PCD proportion = 0.22), *VEGFB* expression was elevated compared to sample S71 (PCD proportion = 0.04) (**Extended Data Fig. 10f**). Hypoxia signature scores were also increased in PCD regions (**Extended Data Fig. 10g,h**), highlighting their angiogenic phenotype.

We next evaluated whether PCD niche abundance predicted response to anti-angiogenic therapy (**Fig. 6g**). In the TCIA.985G-EY35 cohort^54^ (282 WSIs from 76 ovarian and peritoneal serous carcinoma patients treated with bevacizumab), PCD-positive patches were enriched in bevacizumab-sensitive cases, whereas resistant tumors displayed lower PCD scores (**Fig. 6h, Extended Data Fig. 10i, Supplementary Table 11**). Stratification by median predicted PCD score revealed a significantly higher response rate in PCD-enriched samples (p=0.0057; **Fig. 6i**).

Together, these findings demonstrate that the PCD niche is not only a marker of poor prognosis but also a predictive biomarker for bevacizumab response. By leveraging H&E slides, POWER provides a clinically practical framework for risk stratification and therapy selection in HGSOC.

## Discussion

This study represents a comprehensive, large-scale effort integrating Visium HD spatial transcriptomics, multi-omics sequencing, and pathology foundation models to unravel the molecular, cellular, and spatial complexities of gynecologic cancers, including OC, EC, and CC. By linking genomic and transcriptomic alterations with spatial context, we highlight the importance of characterizing shared cellular architectures, TME heterogeneity, and immune infiltration patterns across these malignancies.

We identified conserved cellular compartments across FGTs, yet their varying distributions reflected distinct intratumoral immune infiltration levels and delineated reproducible spatial subtypes. Molecular clusters derived from RNA-seq showed strong concordance with spatial subtypes, underscoring the bidirectional relationship between molecular programs and spatial architecture. While previous studies have mapped transcriptional heterogeneity in OC, EC, and CC^8,12,55–57^, the influence of molecular drivers on tumor spatial organization remains poorly understood. Here, we demonstrate that HPV-induced suppression of TGFβ signaling prevented myCAF accumulation near epithelial regions in CC, thereby reshaping immune infiltration. Conversely, spatial immune exclusion facilitated the proliferative phenotypes of serous tumors within Immune-restricted subtypes. These findings emphasize that tumor architecture is not a passive outcome but an active determinant of tumor phenotypes, shaping proliferation, invasion, and immune evasion^58,59^.

Through integration of WES and spatial transcriptomics, we systematically mapped CIN heterogeneity at both bulk and spatial levels. While serous and endometrioid tumors share Müllerian origins^6,7^, our analyses exhibited distinct genomic landscapes: serous tumors showed *TP53* mutations, extensive CNVs, and *MYC*/*PIK3CA* amplification, whereas endometrioid tumors carried MSI-high status and *ARID1A*/*PTEN* mutations. These differences determined divergent spatial immune phenotypes: CIN-high serous tumors displayed immune-cold states, while endometrioid tumors were immune-hot. Prior research has shown that CIN fosters clonal evolution and immune evasion^16,34,60–63^, but these insights were largely limited to bulk or scRNA-seq analyses. Our study extends this by demonstrating, at single-cell spatial resolution, that CIN-high regions are spatially segregated from immune cells and enriched for MYC/PI3K signaling, providing direct evidence for spatially restricted clonal selection and immune exclusion.

In HGSOC, we uncovered the Proliferating Core Desert (PCD) niche-a malignant spatial domain characterized by CIN-high epithelial cells, elevated MYC/PI3K signaling, metabolic activation, and profound immune exclusion. While prior scRNA-seq studies in HGSOC identified similar transcriptional signatures^64,65^, our spatial analyses reveal how these programs are consolidated within a discrete architectural unit that persists after chemotherapy. Importantly, we show that the PCD niche is not isolated but interacts with specific stromal and immune cell subtypes: CAF-rich boundary niches that remodel ECM via TGFβ signaling, and immunosuppressive EREG+ TAMs that co-localize with CD24/CD47+ epithelial cells. These interactions collectively promote immune evasion and chemoresistance. Consistent with reports that TAMs mediate T-cell exhaustion after chemotherapy^66^ and fibroblasts reduce platinum accumulation in ovarian cancer cells^67^, our findings suggest that CAF and TAM subpopulations act as protective neighbors of the PCD niche, sustaining tumor recurrence. This spatial perspective highlights opportunities for therapies targeting CAF–tumor or TAM–tumor interactions.

Finally, by leveraging the pathology foundation model UNI, we developed POWER, a computational framework that predicts PCD niche abundance from H&E slides. POWER robustly stratified patient prognosis in TCGA-OV and predicted bevacizumab response in an independent clinical cohort. These findings align with emerging multimodal AI approaches in oncology, such as iStar^68^ for predicting spatial gene expression and OmiCLIP^69^ for spatial omics integration. However, most existing frameworks are limited by the resolution and scope of current datasets. Our study provides a large-scale single-cell resolution spatial dataset with paired multi-omics, offering a unique training resource for future AI-driven pathology models.

This work has several implications. First, the PCD niche emerges as a clinically actionable biomarker for both prognosis and therapy selection. Second, our findings argue for a spatially informed view of clonal evolution, where genomic alterations shape not only transcriptional programs but also niche-level architectures and cell–cell interactions. Third, the integrative computational framework demonstrates how spatial omics can be translated into clinically practical workflows via digital pathology. While our dataset spans 100 tumor sections, larger and more diverse cohorts are needed to validate PCD prevalence and therapeutic relevance across populations. Functional validation of ligand–receptor interactions, such as MIF–CD74 and CD24–SIGLEC10, will be critical to establish causality. Finally, while POWER demonstrates strong performance, prospective trials are needed to confirm its utility in guiding therapy decisions.

In conclusion, this study provides the most comprehensive spatial and molecular atlas of gynecologic cancers to date. By integrating high-resolution spatial transcriptomics, multi-omics, and AI pathology, we identify a malignant PCD niche that orchestrates immune exclusion, chemotherapy resistance, and angiogenesis. The development of POWER demonstrates the feasibility of translating spatial biology into clinical tools. Together, these advances provide a foundation for spatially informed precision oncology in gynecologic cancers, with direct implications for biomarker development, therapeutic targeting, and patient care.

## Methods

### Human participants

A cohort of 477 patients with gynecologic tumors was enrolled from the Chinese PLA General Hospital in this study (**Tables S1 and S2**). All patients underwent surgical resection, either before or following systemic treatment with follow-up time ranging from March 2014 to April 2024. During sample collection, necrotic areas were carefully avoided. The majority of tumor specimens were sent to the pathology department, where they were fixed in 10% neutral formalin for histological examination, followed by paraffin embedding for long-term preservation. This allowed for future 10x Visium HD spatial analysis. The remaining tumor tissue was immediately frozen in liquid nitrogen upon excision and stored at −80□°C, while peripheral blood samples (5 mL or at least 1 mL) were collected in EDTA tubes and transported to the laboratory. To minimize processing time, we implemented a standardized workflow that involved collaboration among surgeons, project managers, and laboratory research technicians according to TCGA SOPs (Standard Operating Procedure) (https://brd.nci.nih.gov/brd/sop-compendium/show/701). After independent review by two professional gynecologic pathologists, DNA and RNA were extracted from the same tissue block and then subjected to a comprehensive multi-omics analysis, including whole exome sequencing (WES), RNA sequencing, miRNA sequencing, and whole-genome bisulfite sequencing (WGBS). Patient recruitment and data collection were conducted in accordance with the approved Ethics Committee of Chinese PLA General Hospital #S2022-403-01 after obtaining written informed consent from all participants.

### FFPE tissue preparation for Visium HD spatial gene expression

FFPE tissue samples obtained from PLA General Hospital underwent rigorous quality assessment, including RNA integrity number (RIN) analysis. Only samples with a DV200 value exceeding the quality threshold (>200 nucleotides) were included in the study. Pathologists, clinicians, and scientific researchers collaboratively identified 6.5 x 6.5 mm regions of interest (ROIs) on H&E-stained sections. Following the 10x Genomics Visium HD FFPE Tissue Preparation Handbook (CG000684), 5 μm thick sections were cut, spread out in RNase-free water (Milli-Q) at 42°C, adhered to the slide (Fisher Scientific #1255015), and then air-dried for 30 minutes at room temperature and subsequently for 3 hours at 42°C. The follow-up experiment was conducted after overnight drying at room temperature. After deparaffinization, H&E staining, and imaging, the slides were subjected to the Visium HD assay for spatial gene expression profiling. Library preparation and sequencing were performed by Labway, Shanghai, China.

### DNA and RNA isolation, quantification, and qualification

Genomic DNA was extracted using the Qiagen MinElute Kit, and its purity and concentration were assessed by 1% agarose gel electrophoresis and Qubit 2.0 fluorometer (using the Qubit DNA Assay Kit), respectively.

RNA was isolated using TRIzol reagent and subjected to rigorous quality control. The integrity of the RNA was assessed by 1% agarose gel electrophoresis to exclude degradation and contamination. Additionally, the purity and concentration of the RNA were evaluated using the NanoPhotometer spectrophotometer (IMPLEN) and the Qubit RNA Assay Kit with a Qubit 2.0 Fluorometer (Life Technologies), respectively. Finally, the RNA integrity number (RIN) was determined using the RNA Nano 6000 Assay Kit and the Bioanalyzer 2100 system (Agilent Technologies).

### WES library preparation, exome capture, and sequencing

DNA quality and quantity were assessed using a NanoDrop spectrophotometer and an Agilent 2100 Bioanalyzer. Exome capture was performed using the SureSelectXT Human All Exon V6 Kit (Agilent Technologies, Santa Clara, CA). Briefly, 1 µg of genomic DNA was sheared to an average size of 150-200 bp using a Covaris S220 focused-ultrasonicator. The fragmented DNA was then end-repaired, A-tailed, and adapter-ligated using the KAPA Library Preparation Kit (Kapa Biosystems Inc., Wilmington, MA) according to the manufacturer’s protocol. The adapter-ligated fragments were hybridized to the SureSelectXT Human All Exon V6 probes and captured using streptavidin magnetic beads. After washing, the captured DNA fragments were amplified by PCR to generate a sequencing library. The quality of the constructed libraries was assessed using an Agilent 2100 Bioanalyzer. The concentration of each library was quantified by quantitative PCR (qPCR) using a KAPA Library Quantification Kit (Kapa Biosystems Inc.). Qualified libraries were pooled and sequenced on a NovaSeq 6000 (Illumina) using a 150 bp paired-end sequencing strategy. Sequencing was performed according to the manufacturer’s recommended protocols.

### RNA-seq Library Preparation and Sequencing

The quantity and quality of the extracted RNA were assessed using a Nanodrop One Microvolume UV-Vis Spectrophotometer and Qubit 2.0 Fluorometer with the Qubit RNA HS Assay Kit and an Agilent Bioanalyzer 2100. Subsequently, 10 ng of total RNA was used as input for library preparation using the TruSeq RNA Exome Library Prep Kit (Illumina) following the manufacturer’s protocol. Briefly, mRNA was enriched using poly-T oligo-attached magnetic beads. The enriched mRNA was then fragmented, reverse transcribed into first-strand cDNA, and subsequently converted into double-stranded cDNA. The cDNA fragments were end-repaired, A-tailed, and ligated with Illumina sequencing adapters. Finally, the ligated fragments were PCR amplified with 15 cycles and purified using AMPure XP beads. The constructed RNA-Seq libraries were quantified using Qubit RNA HS Assay Kit and pooled in equimolar amounts. The pooled library was sequenced on an Illumina NovaSeq 6000 using a SP Flow Cell and NovaSeq XP Reagent Kit.

### WGBS Library Preparation and Sequencing

One hundred nanograms of genomic DNA was mixed with 0.5 ng of unmethylated lambda DNA as a spike-in control. The DNA mixture was fragmented to an average size of 200-400 bp using a Covaris S220 focused-ultrasonicator. Bisulfite conversion was performed using the EZ DNA Methylation-Gold™ Kit (Zymo Research) according to the manufacturer’s protocol to convert unmethylated cytosines to uracils. Following bisulfite conversion, Illumina TruSeq adapters were ligated to the fragmented DNA. The adapter-ligated fragments were size-selected using a gel electrophoresis system to enrich for fragments between 200-400 bp. Subsequently, the selected fragments were amplified by PCR to generate a sequencing library. The quality of the constructed libraries was assessed using an Agilent 5400 TapeStation system. The concentration of each library was quantified by quantitative PCR (qPCR) and was required to be greater than 1.5 nM. Qualified libraries were sequenced on an Illumina sequencing platform using a 150 bp paired-end sequencing strategy. Sequencing was performed using Illumina’s sequencing-by-synthesis technology, which involves the incorporation of fluorescently labeled nucleotides during DNA synthesis. The emitted fluorescence signals were captured at each sequencing cycle to determine the nucleotide sequence.

### miRNA-seq Library Preparation and Sequencing

Total RNA was extracted from the tumor sample if available and its integrity and quantity were assessed using the Agilent 2100 Bioanalyzer. 2 μg of total RNA per sample was used as input for library preparation with the NEBNext Multiplex Small RNA Library Prep Set (Illumina). 3’ and 5’ adapters were ligated to the small RNAs, followed by reverse transcription using M-MuLV reverse transcriptase to generate cDNA. The cDNA was then amplified by PCR and purified. The quality and quantity of the resulting sequencing libraries were assessed using Qubit 2.0 and Agilent 2100 Bioanalyzer. Qualified libraries were pooled and sequenced on an Illumina NovaSeq 6000 platform using a 50 bp single-end sequencing strategy. Sequencing data was generated using the sequencing by synthesis principle.

### Visium HD Spatial Gene Expression

FFPE tissue sections were initially placed on plain glass slides and subjected to deparaffinization, followed by ethanol treatments. After ethanol treatment, the sections were stained with hematoxylin and eosin and sealed with glycerol for imaging, as described in the Visium HD FFPE Tissue Preparation Handbook (CG000684). After imaging, the sections were washed and treated with 0.1N hydrochloric acid for decolorization, followed by nucleic acid decrosslinking.

After decrosslinking, the sections were subjected to overnight hybridization with transcriptome probes for 16 to 24 hours, as described in the Visium HD Spatial Gene Expression Reagent Kits User Guide (CG000685). Following hybridization, probe ligation was performed using ligase enzymes, resulting in the formation of stable probe-ligation products. These products were then captured onto 10x chips using the Visium CytAssist instrument. Subsequently, the chips were processed for probe extension, embedding the probes with spatial barcodes essential for accurate spatial location mapping.

Library pre-amplification involved the use of qPCR to assess the quantity of extended probes. Based on the qPCR results, further amplification was performed to increase the abundance of probes containing spatial barcodes. The amplified products were subsequently purified using SPRIselect beads. Following purification, library construction was performed by attaching sequencing adapters and indexes. Subsequently, the sequencing workflow was initiated with the preparation and thawing of reagents in accordance with standard operating procedures (SOPs). Libraries were denatured, diluted, and subsequently sequenced on the NovaSeq X Plus platform with paired-end reads. Libraries were denatured, diluted, and sequenced on the NovaSeq X Plus platform using paired-end reads. Each sequencing cycle, lasting about five minutes, was repeated to ensure high-fidelity results. We performed manual alignment in the Loupe Browser (v.8.0.0) to define and configure the alignment JSON file (https://www.10xgenomics.com/cn/support/software/space-ranger/latest/analysis/inputs/image-cytassist-image-alignment). Subsequently, Space Ranger (v.3.0.0) was employed to map FASTQ files to the human reference genome, align sequencing data with microscope and CytAssist images, and produce gene-barcode matrices for downstream analyses.

### Multiplexed immunofluorescence (mIF)

Slides were deparaffinized in dewaxing solution (Servicebio; 3×10 min), rehydrated in absolute ethanol (3×5 min), and rinsed in distilled water. Then slides were washed in PBS pH 7.4 Endogenous peroxidase was quenched in 3% H₂O₂ for 25 min, followed by PBS washes (3×5 min). For blocking, sections were incubated 30 min with 10% rabbit serum (Servicebio) and 3% BSA (Servicebio). Antibodies were applied and incubated overnight at 4 °C. After PBS washes, HRP-conjugated secondary antibodies were incubated at room temperature for 50 min. Slides were washed in PBS (3×5 min), incubated with the appropriate TSA reagent for 10 min at room temperature, then washed in TBST (Servicebio; 3×5 min). For cyclic staining, residual antibodies were removed with immunostaining antibody stripping buffer (Servicebio): 5 min at room temperature, buffer replaced, then 30 min at 37 °C, followed by TBST washes. After staining cycles, nuclei were counterstained with DAPI for 10 min at room temperature in a light-protected humid chamber (Servicebio). Sections were then washed in PBS, treated with autofluorescence quencher (Servicebio) for 5 min, and rinsed in water for 10 min. After the staining, the images were immediately taken with fluorescence microscope (Nikon).

### DNAscope HD Duplex detection assay

DNAscope HD Duplex detection was performed using the DNAscope™ HD Duplex Reagent Kit (Advanced Cell Diagnostics, Cat #324700) according to the manufacturer’s protocol (Doc. No. UM324700). Adjacent slides were baked at 60°C for 1 hour, then deparaffinized with sequential 5-minute incubations in fresh xylene and rehydrated through two 5-minute incubations in fresh 100% ethanol. Following a 5-minute drying period at 60°C, a hydrophobic barrier was drawn around each tissue section using an ImmEdge pen. Slides underwent RNA removal for 30 minutes at 40°C in a humidity oven, followed by 10-minute incubation with hydrogen peroxide at room temperature, a 15-minute incubation with Protease Plus reagent at 40°C in a humidity oven and heat-induced epitope retrieval using DNAscope Target Retrieval reagent. Hybridization was performed using target-specific DNAscope probes (DNAscope™ Probe-DS-Hs-MYC-O1, ACD, Cat #1123771-C1; DNAscope™ Probe-DS-Hs-CEP8q-C2, ACD, Cat # 1107621-C2) for 15-18 hours at 40°C in the HybEZ II hybridization system. Signal amplification was achieved with the DNAscope HD Duplex amplification reagents, and probes were visualized using DNAscope Fast Red and DNAscope Blue detection reagents, according to the manufacturer’s recommendations. Images were acquired using a VS200 slide scanner and analyzed with the HALO system.

### WES data processing

Raw FASTQ files were preprocessed using fastp (v.0.23.4) with default parameters to remove low-quality reads and adapters. Reads were then aligned to the GRCh38 human reference genome (GRCh38.d1.vd1) using BWA-mem (v.0.7.17) with default settings. Mapping quality was assessed using samtools (v.1.9; https://github.com/samtools/samtools) to ensure high-quality alignments.

### Somatic mutation calling using WES

Somatic mutations, including single-nucleotide variants (SNVs) and small insertions and deletions (indels), were called from each tumor–control pair using the GPU-accelerated somatic pipeline provided by the NVIDIA Parabricks toolkit (v.4.3.0; with parameters: --bwa-options="-Y" and all other parameters set to default; https://www.nvidia.com/en-us/clara/genomics/).

Variants output by the somatic pipeline were annotated with VEP (v.102.0; corresponding to GENCODE v.35) by vcf2maf (v.1.6.20; https://github.com/mskcc/vcf2maf). High-confidence ensemble somatic short variants (depth of tumor□ > □25, depth of control□>□5, variant allele reads count of tumor□>□25, variant allele reads count of control□<□3) were selected for downstream d*N*/d*S* analysis. Furthermore, only variants with “IMPACT” classified as “HIGH” or “MODERATE” were retained, filtering out mutations that did not alter the amino acid sequence in coding regions. The remaining variants were considered significant somatic mutation sites.

### CNV calling using WES

CNVs in tumor samples were called using the R package FACETS^70^ (v.0.6.1) and the R package Sequenza^71^ (v.2.1.2). FACETS analysis was performed using aligned sequence BAM files of tumor and paired normal samples as input, and SNPs were filtered based on mapping quality (≥15) and base quality (≥20). Then, the purity model was applied to estimate the overall segmented copy-number profile, sample purity, and ploidy. Utilizing the dipLogR value inferred from diploid state in the purity model enabled the high-sensitivity model to detect more focal events. Subsequently, the processed data, output as a cncf file, was used to infer CNV segments. For each segmented data line, the total copy number (tcn) was extracted and evaluated against predefined thresholds (gain_cutoff = 3, loss_cutoff = 1.5) to classify segments as GAIN (tcn ≥gain_cutoff) or LOSS (tcn ≤loss_cutoff).

For Sequenza, BAM files of tumor and paired normal samples were used as input and were converted into SEQZ files using the bam2seqz function. The resulting files were normalized using the GC content bias and data ratio. Then, the seqz_binning function was used to bin chromosomal segments into 50 kb. Finally, to acquire segmented copy numbers and estimate cellularity and ploidy, the following parameters were used: min.reads□=□10, min.base.freq□=□0.2, max.reads.count□=□300 and all other parameters set to default using sequenza.extract function.

To obtain high-confidence CNV segments, overlapping regions between the CNV results generated by FACETS and Sequenza were filtered. For these overlapping regions, the more reliable total_cn and status were selected based on segment length. Subsequently, regions longer than 100 kb were retained, and the overlapping segments were integrated into a high-confidence CNV dataset for each sample. Next, bedtools (v.2.31.1) was used to annotate the resulting BED files with the GTF file of GRCh38 (GENCODE v.32).

### CIN score estimation

The CIN score for each sample was calculated as the proportion of chromosomal segments with CNVs, determined by dividing the total length of CNV segments by the total length of the 22 autosomes.

### Tumor purity estimation

The BAM file of tumor and paired normal samples was employed as input data, and FACETS was executed to estimate purity, alongside calling CNVs. The purity model estimated the overall sample purity and ploidy with the following parameters: cval = 150 and maxiter = 1000.

### MSI detection using WES

The MSI status of samples was determined by MSIsensor2 (v.0.1; https://github.com/niu-lab/msisensor2), a tool that utilizes a tumor-control paired module with default parameters. MSIsensor2 calculates the MSI score as the proportion of MSI-positive sites relative to the total number of valid sites. Following the recommended threshold score of 3.5 for MSI^72^, samples with an MSI score ≥ 3.5 were classified as MSI-high, whereas those with an MSI score < 3.5 were categorized as MSI-low.

### Mutational signature analysis

De novo mutational signatures were extracted from 80 CC samples with available paired normal samples. Analyses of mutational signatures were performed by SigProfilerExtraction^73^ (v.1.1.24) with the parameters ‘reference_genome = "GRCh38", opportunity_genome = "GRCh38", minimum_signatures = 1, maximum_signatures = 20, nmf_replicates = 100, cpu = 24, gpu = True, cosmic_version = "3.4"’. The result files under the folder SBS96/Suggested_solution were used for downstream statistics and analysis.

### Calculation of co-occurrence frequency of mutation signatures

First, the total occurrence count (*N_i_* and *N_j_*) for any two distinct mutation signatures (*i* and *j*) was determined individually. Next, the co-occurrence count (*N_ij_* ), representing the number of samples where both signatures were observed simultaneously, was calculated. The normalized co-occurrence value ( *C_ij_* ) of signature *i* and *j* was described:

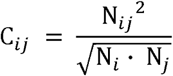

Additionally, for self-pairing of the same signature, the value was set to 1, facilitating subsequent analyses.

### Bulk RNA sequencing processing

Paired-end reads were aligned to the human genome using STAR (v.2.6.1; with parameters: --runThreadN 16, --chimSegmentMin 12, --chimJunctionOverhangMin 12, --chimMultimapScoreRange 10, --chimMultimapNmax 10, -- chimNonchimScoreDropMin 10, --peOverlapNbasesMin 12, --peOverlapMMp 0.1, -- alignSJstitchMismatchNmax 5 -1 5 5, --twopassMode Basic, --quantMode TranscriptomeSAM, and all other parameters set to default), and the resulting BAM files were sorted, indexed, and analyzed for key metrics using samtools. Alignment and subsequent gene counts were performed using the genome annotation version GENCODE v.22.

### Gene Expression Quantification using RNA-seq

Gene counts were quantified using Salmon^74^ (v.0.13.1; with all parameters set to default). To convert transcript-level counts into gene-level counts from SF files and integrate the expression data across multiple samples, the R package tximport^75^ (v.1.28.0) was employed. TPM values of the whole transcriptome were quantified and exported using the tximport package. Downstream analyses were restricted to protein-coding genes, as defined by the BioMart dataset (Homo sapiens genes, GRCh38.p14; https://www.ensembl.org/biomart/martview) with the filter set to “Gene type: Protein coding”.

### Molecular subtype clustering using RNA-seq

To identify distinct molecular subtypes, NMF consensus clustering was performed on normalized TPM values of 250 OC, 124 EC, and 84 CC samples. The R package ConsensusClusterPlus^76^ (v.1.66.0) was used to determine the optimal number of clusters with the following parameters: ‘maxK=10, reps=1000, pItem=0.8, pFeature=1, clusterAlg=’hc’, distance=’pearson’’. Clustering with k=6 gave a stable k-factor decomposition and sample by sample correlation matrices.

#### TME analysis using RNA-seq

To predict the immunosuppressive level using RNA-seq datasets, Tumor Immune Dysfunction and Exclusion (TIDE)^77^ analysis was employed using the tidepy tool on TPM-normalized gene counts to estimate the relative gene signature enrichment scores of IFN-γ signaling (IFNG), cytotoxic T lymphocyte (CTL), and CAF in each sample. To determine the abundances of different cell types in FGT, R package CIBERSORT^21^ (v.1.0.4) was used to estimate the relative cell composition in each sample. The parameters of CIBERSORT function were set as ‘perm = 100, QN = "FALSE", absolute = "TRUE", abs_method = "sig.score"’ and all others were set to default.

### WGBS data processing

WGBS data of FGT samples were analyzed using the nf-core methylseq pipeline^78^ (v.2.4.0; https://nf-co.re/methylseq). The pipeline is built using Nextflow (v.23.04.1; https://github.com/nextflow-io/nextflow) and integrates pre-processing, genome alignment, post-processing, and final quality control stages. The parameter “--aligner bwameth” was used, and GRCh38 human reference genome served as a reference.

### Extracting and Filtering CpG sites

To extract and filter CpG sites from BED files, the R package methylKit^79^ (v.1.26.0) was used, converting BED files into tables. CpG methylation was estimated using the ’methRead’ function in the MethylKit package with the parameter mincov = 5.

### Methylation score calculation for promoters

To estimate gene promoter level methylations, RefSeq transcription start site (TSS) information for hg38 genome build was obtained from the UCSC genome browser. A fixed window size of 1,000 bp upstream of the TSS was employed for each gene and the methylation score (β value) for each gene was calculated by dividing the total number of methylated CpG sites by the total coverage of CpG sites within this region in each sample.

### Differentially methylated region analysis

To compare methylation levels at gene promoters between serous and endometrioid subtypes, we merged CpG site annotations filtered to regions within 1,000 bp upstream of TSS across serous and endometrioid samples. Methylation data were processed using MethylKit’s ’methRead’ function with a minimum coverage threshold of 5 reads to ensure reliable quantification. The merged data was analyzed for differentially methylated region identification using MethylKit’s ’calculateDiffMeth’ function, which applies a chi-squared test and adjusts for q-values.

### miRNA-seq data processing

FASTQ files were preprocessed using fastp to remove adapters and filter low-quality reads with default settings. The miRBase v22 mature database (http://www.mirbase.org/) was used as the reference for miRNA transcripts, and the index for this reference database was generated using build tool from bowtie (v.1.3.1). Sequenced reads were aligned to the reference database using bowtie with parameters ‘-n 0 -m 1 --best --strata’.

To quantify miRNA transcript counts, the output from the alignment step was first sorted using the sort tool from samtools. Subsequently, indices for the sorted BAM files were generated using samtools index, and aligned miRNA transcript counts were calculated using samtools ’idxstats’ tool.

### Differential expression analysis using miRNA-seq

Differentially expressed miRNA transcripts among the four molecular subtypes defined by RNA-seq data were detected by DESeq2 (v.1.42.0). P-values were adjusted for multiple testing using the Benjamini–Hochberg procedure.

### Visium HD data quality control

Squidpy Python package (v.1.2.3) was used to import the gene-barcode matrices produced by the Spaceranger count pipeline. For each sample, only the first occurrence of each gene name was retained, by removing duplicates with identical names but differing ENSG IDs. Additionally, IGKC and IGHG1 were deleted due to their abnormal spatial diffusion, which could affect unrelated adjacent regions. Besides, genes detected in fewer than five bins and bins with fewer than five total detected gene counts were removed to ensure data quality and analytical reliability.

### Visium HD data normalization

To eliminate sequencing depth differences among bins, gene expression values were normalized for each bin according to:

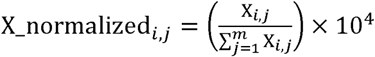

where *X_i,j_* is the count value of the *j*-th gene in the *i*-th bin, *m* is the total number of genes in each sample.

Then a logarithmic transformation was performed to scale the dynamic range of the data. Expression levels (*E_i,j_*) were quantified as shown by:

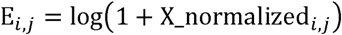

### Visium HD spatial distance quantification

The location of each bin was determined based on its specific row and column coordinates within the probe array. To quantify distances between cell-cell pairs, cell-cell group pairs, and cell group-cell group pairs, and measure the interaction probability in ST data, three spatial distance metrics were defined: physical distance, neighborhood cell type proportion, and contact likelihood score. Physical distances were determined by calculating the Euclidean distance between the bin coordinates of each cell-cell pair.

The contact likelihood score was derived from pairwise contact frequencies^80^. For each cell group-cell group pair (e.g., distinct CNV states or cell types), the contact frequencies were defined as the observed number of contact edges between groups across all neighborhoods, with the contact edges determined from the spatial network constructed within each neighborhood. Then, contact likelihood score (*S_i,j_*) between group i and group j was calculated according to the following formulas:

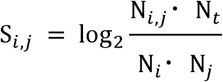

where *N_i,j_* is the number of contact edges between cells in group *i* and group *j*, and *N_t_* is the total number of contact edges (*∑_i,j_ N_i,j_* ), while *N_i_ = ∑_j_ N_i,j_* and *N_j_* = *^∑^_i_ ^N^_i,j_*.

### Visium HD cell type annotation

#### Quality control

All processes in the cell annotation section were performed individually for each sample. Using the preprocessed gene expression matrix after filtering specific genes, 2 μm spots with fewer than one detected gene or fewer than five UMIs were excluded. Similarly, 8 μm bins with fewer than three detected genes were removed. Additionally, genes detected in fewer than five 2 μm spots were filtered out.

#### Normalization

The raw count matrix for the 2 μm spots was normalized to a target count of 10,000 per spot using the normalize_total function from the Scanpy package, followed by a log transformation (log1p) for downstream analysis.

The relative abundance of each cell type was first calculated based on predefined marker genes for different cell types. For cell type *j*, its relative abundance *WCT_j_* was calculated as:

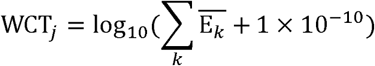

where *k* denotes the marker gene *k* from cell type *j*, and 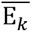 denotes the average expression of gene *k* in spots that express gene *k*.

Next, the number of expressed marker genes for cell type 1 in spot, was calculated as:

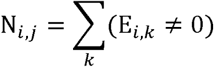

where *E_i,k_* denotes the expression of gene *k* in spot *i*.

The probability score of a spot *i* belonging to cell type *j* was then computed as:

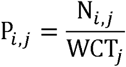

Finally, each 2 μm spot was assigned to the cell type with the highest probability score if that score is not zero. Otherwise, the spot was labeled as "Others".

#### Cell type assignment

The spatial distribution of cell types was determined by mapping annotations from high-resolution 2 µm spots to lower-resolution 8 µm bins. For each cell type within a sample, the weight was calculated as the proportion of annotated 2 µm spots corresponding to that cell type relative to the total number of annotated spots on the slide. The weight of cell type *j* was calculated as:

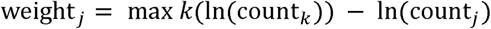

where *k* represents any given cell type.

In almost all samples, the cell type ’Others’ was the most prevalent, providing a baseline for standardizing the weights of other cell types relative to the ’Others’ count. For one outlier sample (S92), where this pattern did not hold, weights were adjusted to avoid the over-classification of biologically relevant cell types as ’Others’.

Next, 8 µm bins were annotated based on the pre-annotated 2 µm spots within them. An 8 μm bin was assigned the label “Others” only if all corresponding 2 μm spots within the bin were categorized as “Others”. Otherwise, for each 8 μm bin *t*, the weighted cell type count for cell type *j* was calculated as:

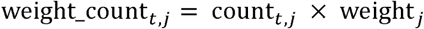

where count*_t,j_* represents the number of 2 μm spots labeled as cell type *j* within the corresponding 8 μm bin *t*.

Finally, each 8 μm bin was annotated as the cell type with the highest weighted count. To minimize the impact of cell type mixing, within each 8 μm bin, only 2 μm spots classified as the same cell type as the 8 μm bin or labeled as “Others” were retained. The gene expression data from these selected 2 μm spots were then aggregated to reconstruct the expression matrix for the 8 μm bins.

### Identification of cellular compartment using Visium HD data

To identify conserved spatial structures in FGTs through their spatial neighborhood profiles, we applied a clustering approach based on neighboring celltype composition. We first defined the neighborhood region of each cell. Specifically, each cell was designated as the center, and its neighborhood was defined by four concentric layers, with the outer layer encompassing a square area of 72 μm side length. Then a single-cell neighborhood matrix was generated through quantification of local cellular composition of each cell (62,428,567 cells × 11 celltypes), with the rows encoding celltype proportions within the spatial neighborhood of individual cells. To systematically characterize celltype composition patterns and reduce computational complexity during clustering, we applied NMF to the single-cell neighborhood matrix. This approach enabled the identification and quantification of localized cellular features, obtaining a cell-by-feature matrix (W matrix) and a feature-by-celltype matrix (H matrix). We employed the ’NMF’ function from ’sklearn.decomposition’ package and set the parameters as ’ n_components=5, init=’random’, random_state=42 ’. Subsequently, the W matrix was used to perform cell clustering using a Gaussian Mixture Model (GMM) implemented in CellCharter. To identify the optimal cluster count yielding a stable solution, the GMM was executed five times for each candidate number within the range of 5–20 clusters. Ultimately, 14 clusters were selected as the optimal solution for CellCharter-defined spatial clusters across 100 Visium HD samples, demonstrating maximal stability in clustering consistency. Furthermore, these spatial clusters were manually annotated based on their celltype compositions and integrated into 11 cellular compartments with shared compositional profiles.

### Differential gene expression analyses using Visium HD data

Analyses of differential gene expression between groups of cells were performed using the rank_genes_groups function from scanpy library. To normalize expression levels, cells with fewer than 10 expressed genes and genes expressed in fewer than 3 cells were excluded based on the raw count matrix. Subsequently, raw counts were normalized per cell, and a log(count+1)-transformed matrix was generated for downstream DEG analyses.

### Pathway and signature analyses using Visium HD data

Gene Set Enrichment Analysis (GSEA) was performed using the R package clusterprofiler (v.4.10.0). The normalized enrichment score (NES) and p-value based on GOBP, HALLMARK, KEGG, and REACTOME gene sets were determined by GSEA with 10,000 random permutations of gene labels.

The signature scores of cells were calculated for each gene sets using the score_genes function from Scanpy, applied individually to each sample. These scores represented the relative activity of the specified gene signatures in each cell. To enhance spatial visualization, a smoothing process was applied to the signature scores: for each cell, the score was recalculated as a weighted sum of its own score and those of its 8 nearest neighbors.

### Cell-cell communication analysis

CellPhoneDB^81^ (v.2.0) was applied to infer the ligand-receptor interaction between cell types within the OPSCC scRNA-seq dataset. The interaction partners were computationally identified from the dataset and systematically referenced with curated ligand-receptor interaction databases by the R package CellChat^82^ (v.1.6.1). Intercellular ligand-receptor interactions were characterized at single-cell resolution through: (1) mean expression by cell subtypes, (2) Monte Carlo permutation testing (p<0.05 FDR-corrected), and (3) cellular detection filters excluding ligand/receptor genes observed in <10 cells per subtype.

### CNV inference for Visium HD data

The infercnvpy Python package^31^ (v.0.4.5; https://github.com/icbi-lab/infercnvpy) was employed to infer somatic CNVs from Visium HD data. Infercnvpy was run at the sample level and only with post-quality control filtered data. The location of each gene on the chromosome was annotated according to the human reference GRCh38-2020-A. The 17,332 genes on 22 autosomes were included in our subsequent analysis.

Recognizing the sparsity of HD data and the fact that the mean diameter of epithelial cells exceeds 8 μm^83^, a spatial smoothing step was implemented to average the raw expression of neighboring epithelial cells, enhancing the signal (counts) while preserving the spatial context. For each epithelial cell, if any epithelial cells were present among its eight nearest neighboring cells, the gene expression was smoothed by computing a weighted sum of its own gene expression and the averaged gene expression of epithelial cells among its eight nearest neighboring cells, as follows:

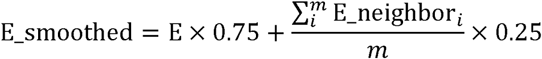

where E is the original expression data of the central epithelial cell, and E_neighbor represents the original expression data of each epithelial cell among its eight nearest neighbors. The variable *m* denotes the number of epithelial cells present among the eight neighboring cells.

Then normalization and log transformation were performed as described in the “Visium HD data quantification and processing” part. After preprocessing the gene expression data, the infercnvpy package was used for CNV inference. Non-epithelial cells, excluding B cells (due to their high expression of specific IG genes), were selected as the reference population, serving as a normal baseline for detecting CNVs in epithelial cells from the same sample. Then, using parameters window_size=190, step=1, and other default parameters, CNV profiles, and corresponding heatmaps were generated to display inferred copy number alterations for all genes in each cell, enabling a comprehensive visualization of genomic variability across the dataset.

### CNV profile clustering and scoring on Visium HD data

PCA dimensionality reduction (n_components=20) and Leiden clustering (with n_neighbors=15, resolution=0.6, and n_iterations=2) were performed on the inferred CNV profiles of each sample’s epithelial cells to identify distinct CNV clusters.

The CNV score was computed for each cell by averaging the absolute values of the inferred copy number alterations across all genes as:

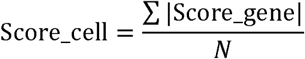

where Score_gene is the inferred copy number alterations of each gene and *N* is the number of genes in the cell.

#### CNV level stratification

Based on the CNV scores, we stratified all cells within a sample into four distinct levels using the quartile method. Specifically, cells with CNV scores between the third to fourth quartile were classified as the top level. Cells with scores ranging from the median to the third quartile were assigned to the high level. Cells with scores ranging from the first quartile to the median were categorized as the low level, while cells with scores below the first quartile were designated as the bottom level.

#### CNV state classification

For each sample, the CNV score vector for each Leiden cluster was computed by averaging the inferred copy number score for each gene across all cells within the

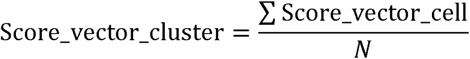

, where Score_vector_cell is the CNV score vector which contains the CNV score of all genes in a cell and *N* is the number of cells in the cluster.

Subsequently, CNV clusters of all samples were divided into 5 CNV states (Chr8q-amp, Chr20-amp, Chr18-amp, CIN-moderate, and CIN-low) by performing hierarchical clustering on their Score_vector_cluster with parameters method=’ward’ and distance_threshold = 2.4.

#### CNV profile similarity analysis

The CNV score vector from HD data for each sample was determined by averaging the gene scores of the inferred copy number alterations across all cells within the sample, i.e., Score_vector_sample. Based on this, pairwise cosine similarities were calculated between CNV profiles inferred from Visium HD data and WES data, respectively. A total of 17,197 genes shared between the HD and WES datasets were identified by computing their intersection. Then we calculated cosine similarity of a sample as follows:

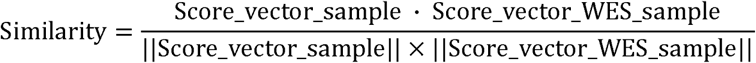

where Score_vector_WES_sample is the WES-based CNV profile of the same sample in HD data.

### Spatial clustering on 40 HGSOC samples

Spatial clustering groups 8 µm bins by integrating the gene expression profiles of each bin with the features of its neighboring bins, capturing both local and contextual spatial information. In this study, CellCharter (v.0.2.2; https://github.com/CSOgroup/cellcharter) performed spatial clustering on the integrated HD data from 40 HGSOC samples through the following steps: (1) constructing a spatial network based on the coordinates of each bin; (2) performing dimensionality reduction and batch effect correction; (3) aggregating the features of each bin with those of its spatial neighbors; and (4) clustering the aggregated feature matrix across all bins.

#### Spatial network construction

Given that 8 µm bins are arranged in continuous arrays, each bin was represented as a node, and edges were established between directly adjacent nodes. Specifically, functions in the Squidpy package were implemented to construct the network for each sample by assigning the four nearest surrounding bins (top, bottom, left, and right) as neighbors for connectivity.

##### Dimensionality reduction and batch effect removal

scVI^84^ was used to perform dimensionality reduction and batch effect removal on the HD data from 40 samples. scVI is built on a variational autoencoder (VAE) architecture, designed to learn low-dimensional embeddings that can effectively reconstruct the original input data. The raw count matrix was used as the input for scVI, with the batch key set to the sample ID. During the training process, the maximum number of epochs was set to 5, and GPU acceleration was employed to enhance computational efficiency. Finally, 10-dimensional latent representations were generated for a total of 23,100,945 bins from 40 HGSOC samples.

#### Neighborhood feature aggregation

CellCharter defines the *l*-neighborhood of a bin as the collection of bins that are at most *l* steps away from the bin within the constructed network. In this study, *l* = 3 was set, and for each bin, its feature vector was concatenated with *l* layer vectors, where each vector represented the average features of bins located *i*-steps away within the network, for *i* ∈ [1, *l*].

#### Spatial clustering

All the bins were clustered based on aggregated features using a GMM implemented in CellCharter. Furthermore, to determine the optimal number of clusters yielding a stable solution, the GMM was run five times for each candidate cluster number within the range of 6 to 24. At last, 21 clusters were selected as the optimal number for CellCharter-defined clustering across the 40 HGSOC samples, achieving the highest stability.

### Macrophage subtype decomposition in Visium HD data

To refine macrophage subtype annotations in our Visium HD datasets, we used the public single cell transcriptomics dataset of human HGSOC as a reference^52^, and applied Spotiphy (https://github.com/jyyulab/Spotiphy), a computational toolkit enabling spatially resolved celltype deconvolution, to map celltype annotation information from scRNA-seq to ST data.

First, Spotiphy performed feature selection for TAM subtype-specific informative genes, deriving comprehensive signature reference profiles from scRNA-seq data. This step was executed using the ’spotiphy.sc_reference.marker_selection’ function with parameters: n_select=50, threshold_p=0.1, threshold_fold=1.0, q=0.15. Following this, we obtained subtype-specific gene signatures containing up to 50 genes per TAM subtype, which were applied to deconvolute individual macrophage bins from Visium HD datasets using ’spotiphy.deconvolution.estimation_proportion’ function. After deconvolution, a cell-by-subtype matrix was generated, where each row quantifies the proportion of each subtype within a macrophage bin. Finally, we established refined subtype assignments by selecting the subtype with the maximal value in each macrophage bin if no values were lower than 0.3; otherwise, the macrophage was classified as ’Undefined’.

### Construction of POWER (Pathology-based Ovarian cancer-specific Workflow for Evaluating Risk)

#### In-house WSIs processing

Whole slide images (WSIs) from 40 HGSOC patients with associated clinical information were obtained. WSIs were segmented into 6.5 × 6.5 mm^2^ images to align with the spatial capture areas of the Visium HD platform. Each segmented image was further cropped into 1.6×1.6 mm² patches. Using the spatial coordinates and their correspondence to pixels in the high-resolution H&E image, the spatial distribution of gene expression features for the respective regions was obtained. Patches with fewer than 3,000 cells were treated as non-specific background patches and filtered out. A total of 627 cropped patches were obtained, with a mean resolution of 6393×6393Lpixels at ×40 magnification. All image cropping was performed using the Python package pyvips (v.2.2.1; https://github.com/libvips/pyvips).

#### Feature extraction

For feature extraction, all patches were resized to 224□× 224□pixels and normalized using the mean and standard deviation parameters of ImageNet (https://image-net.org/). Feature embeddings for each patch were generated independently using a frozen pretrained image encoder, UNI. The UNI encoder’s backbone followed the ViT-L/16 architecture, a large Vision Transformer (ViT) model with 24 transformer layers, 16 attention heads, and an embedding dimension of 1024.

Each 224 × 224 patch was divided into 16 × 16 pixel sub-patches, forming a 14 × 14 grid of sub-patch embeddings through a 2D convolutional layer with a kernel size and stride of 16. Each grid cell contained a 1024-dimensional vector. The resulting 2D grid was flattened and transposed into a [196 × 1024]-dimensional sequence of embeddings, suitable for transformer-based attention. Finally, the UNI encoder produced a 1024-dimensional feature vector for each patch.

#### Calculation of PCD niche proportion

For each patch, the corresponding PCD niche proportion was calculated. The spatial clustering results were mapped to each cropped patch obtained from the H&E images using the spatial coordinates from the Visium HD data. Each 8 μm bin has a corresponding niche label, and the number of PCD niche bins within each patch was counted to represent the area corresponding to the PCD niche region. Similarly, the total number of bins detected in each patch was counted as the area of the tissue region. Then, the proportion of PCD niche of a patch is

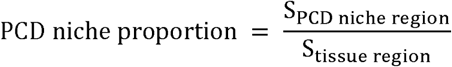

Finally, an in-house dataset with dimensions of 627×1024 and paired PCD niche proportion labels was constructed, where 627 represents the total number of patches obtained from the H&E images, and 1024 refers to the embedding dimension of each patch encoded by UNI, with spatial clustering information incorporated for each patch.

#### Prediction model construction

For each patch, a PCD niche status was assigned as “true” if the PCD niche proportion exceeded 20%. During model selection, four models-MLP, SVM (RBF), SVM (Linear), and Random Forest-were evaluated. Based on the area under the ROC curve and the distribution of predicted PCD niche proportions, the MLP model was selected as the prediction model.

The splitting ratio was 6:2:2 for the training, validation, and test set. After hyperparameter tuning, the final model architecture comprised three fully connected layers with an input dimension of 1024, intermediate layers of 256 and 64 dimensions, and a single output representing the predicted proportion of PCD niche. ReLU activation functions were applied to the hidden layers, and the model was trained using Mean Squared Error (MSE) as the loss function for 200 epochs on the training set, with an Adam optimizer and a learning rate of 1×10^−3^. Pearson’s correlation coefficient was calculated between the predicted and true proportions on the test set to assess the prediction accuracy.

#### Survival analysis based on the TCGA-OV cohort

H&E-stained image data and corresponding clinical data were available for 83 patients within the TCGA-OV cohort. To ensure that each patch had a similar microns-per-pixel scale to the training set, the WSIs (n = 83) were divided into contiguous 5000 × 5000 pixel patches at ×40 magnification. Patches with an average RGB value below 240 were considered non-specific background patches and filtered out. The mean number of patches extracted per slide was 207.

As described above, each patch yielded a 1024-dimensional feature vector encoded by UNI. These feature vectors served as input to the MLP model to obtain the predicted proportion for each patch.

A PCD niche-positive patch was defined as a patch where the predicted PCD niche proportion exceeded 20%. This threshold of 20% was chosen based on its ability to maximize the difference between the true positive rate and the false positive rate when binarizing the actual and predicted proportions. Subsequently, a PCD niche score for each whole slide image was calculated as follows:

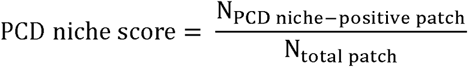

Patients were then stratified based on the median PCD niche score. The survival difference between the groups was compared using a Kaplan-Meier analysis and log-rank test, estimated using the survival R package.

#### Bevacizumab response analysis based on the TCIA.985G-EY35 cohort

After removing slides with missing metadata, 282 slides from 76 patients (68 EOC and 8 PSPC) were collected, sourced from the TCIA.985G-EY35 dataset^54^. The bevacizumab treatment was effective in 160 slides and invalid in 122 slides within the dataset. The WSIs (n = 282) were divided into contiguous 3000 × 3000 pixel patches at ×20 magnification. Patches with an average RGB value below 230 were filtered out to remove background patches, as the background colors vary between different datasets. The mean number of patches extracted per slide is 172.

Using similar strategies, patients were stratified into two groups: predicted PCD niche-enriched and predicted PCD niche-deficient. The bevacizumab response rate difference between the groups was compared using a chi-squared test.

### Statistical analysis

All data analyses were conducted in R and Python environments. Significance was determined using appropriate statistical tests, including the Wilcoxon rank-sum test, Wilcoxon signed-rank Test, ANOVA test, Chi-squared test, permutation test, Kruskal-Wallis sum-rank test, and Pearson correlation test. Survival analysis was performed with the R package survival (v.3.5.7) using the log-rank test. P-values□<=□0.05 were considered statistically significant. Detailed descriptions of the statistical tests are provided in the figure legends and the relevant methods sections.

## Data availability

The raw sequencing data and processed data of multi-omics and Visium HD will be released following publication of the paper. Visualization of the multi-omics data is accessible on the online website (https://stage.pku-genomics.org/). H&E image and CytAssist image files of samples from Visium HD datasets could be downloaded from this website after publication. Single-cell datasets reanalyzed are available through the GEO with accession numbers GSE182227 (OPSCC) and in Mendeley Data (https://doi.org/10.17632/rc47y6m9mp.1) (HGSOC).

## Code availability

The analysis codes are deposited on GitHub (https://github.com/zenglab-pku/FGT-HD).

## Acknowledgements

This work was supported by the National Natural Science Foundation of China (92374116, 32470664, T2321001 to Zexian Zeng; 82404094 to Zhe Zhang), Beijing Natural Science Foundation (L248043, Zexian Zeng), Noncommunicable Chronic Diseases-National Science and Technology Major Project (2024ZD0520600, Zexian Zeng), Sichuan Science and Technology Program (2024YFFK0064, Zexian Zeng), Beijing Advanced Center of Cellular Homeostasis and Aging-Related Diseases (Zexian Zeng), and Peking-Tsinghua Center for Life Sciences (Zexian Zeng). We thank the Optical Imaging Core Facility for experimental platforms. Part of the analysis was performed on the High-Performance Computing Platform of the Center for Life Sciences at Peking University.

## Contributions

T.X., Zhe Zhang, Z.Zeng, and Y.M. conceptualized and designed the study. L.Z., W.Y., M.Y., Zhe Zhang and Y.M. collected the clinical samples. T.X., S.L., and Y.W. conducted bioinformatic analyses under the supervision of Zhe Zhang and Z.Zeng. T.X., S.L., Y.W., Y.H., Zongxu Zhang, Y.Z., and P.Z. constructed the pipeline to analyze the HD data. T.X. and P.Z. built the online database. T.X., S.L., Y.W., Zhe Zhang, and Z.Zeng wrote the manuscript. Y.H., Zongxu Zhang, Y.Z., R.Z., L.W., P.R., C.L. and Y.M. provided edits to the manuscript. Zhe Zhang, Z.Zeng, and Y.M. oversaw the ethical guidelines and data regulation. Z.Zeng supervised the project.

## Corresponding authors

Correspondence to Zhe Zhang, Zexian Zeng, and Yuanguang Meng.

---

**Extended Data Fig. 1:**
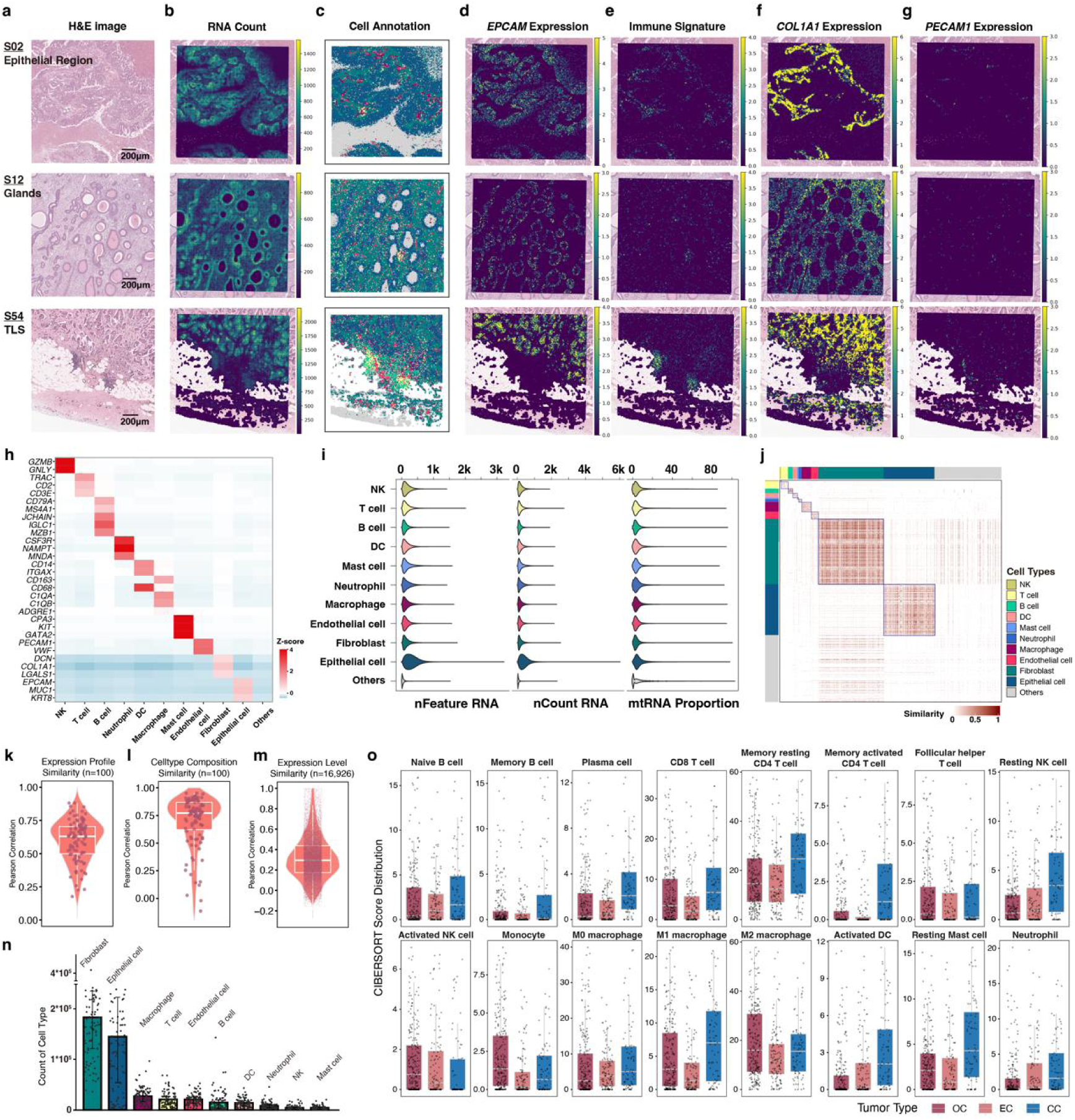
Spatial transcriptomics quality and cell type distributions in the FGT cohort. **a-g,** Representative ROIs from different tumor samples showing H&E-stained images (**a**), RNA counts (**b**), cell type annotations (**c**), *EPCAM* expression (**d**), immune cell markers (**e**), *COL1A1* expression (**f**), and *PECAM1* expression (**g**). All at 8 µm resolution. **h**, Heatmap showing normalized expression Z-scores for marker genes across annotated cell types. **i,** Violin plot of RNA quality scores by cell type. **j,** Cosine similarity of marker-gene expression among cell types, computed from 80,000 randomly sampled cells across OC, EC, and CC cohorts. **k-m**, Validation of Visium HD data against matched bulk RNA-seq data. Pearson correlation coefficients (PCCs) for whole transcriptomic profiles (**k**), deconvoluted cell-type proportions (**l**), and expression similarity across 16,926 shared genes (**m**). **n,** Bar plot of cell-type distributions across 100 samples. **o,** CIBERSORT-derived immune infiltration scores from bulk RNA-Seq data across OC, EC, and CC cohorts.

**Extended Data Fig. 2:**
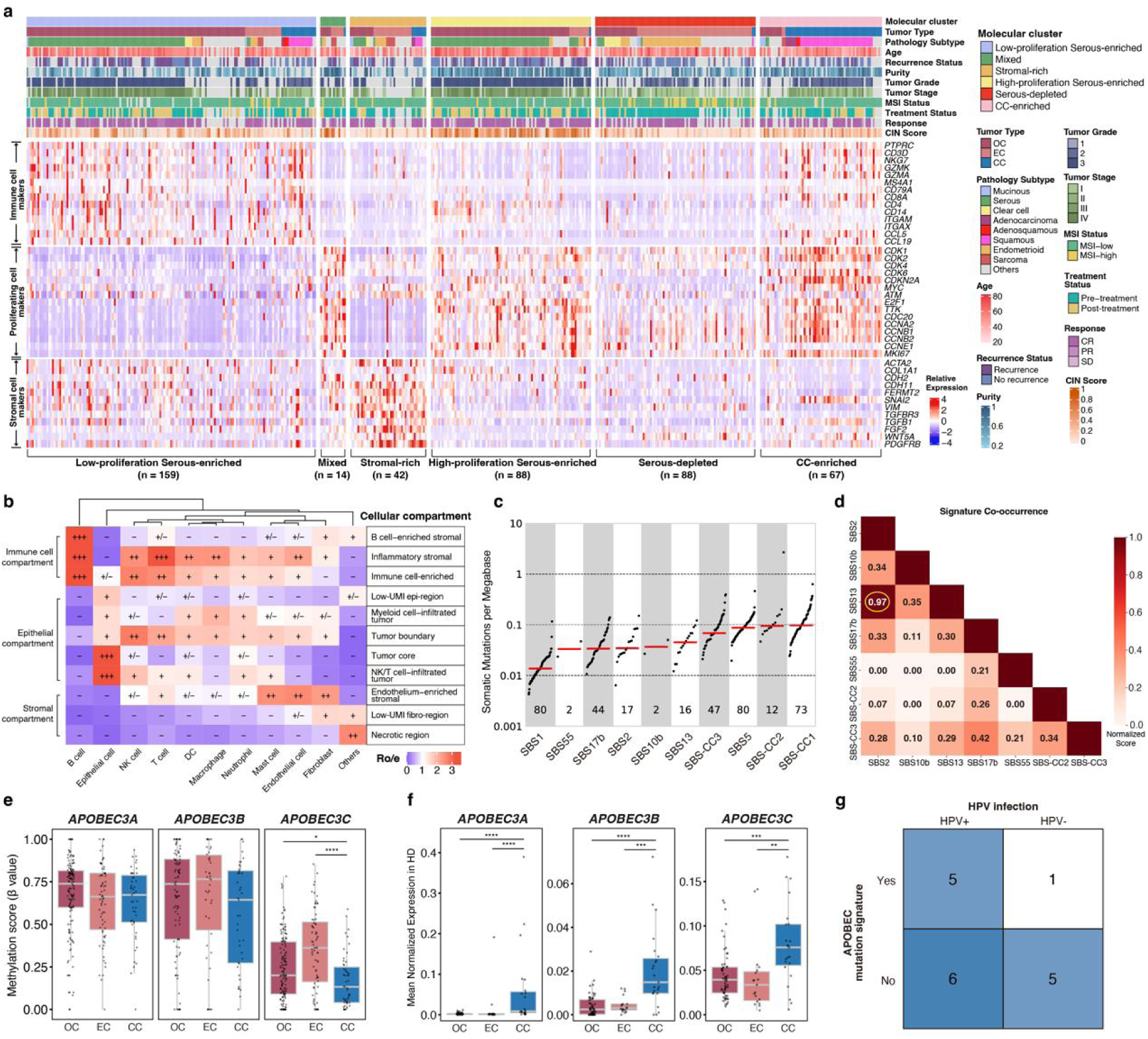
Molecular subtype clustering and characterization of HPV-associated features. **a,** Heatmap of 458 FGT samples clustered into six molecular subtypes using consensus NMF of RNA-Seq data. Immune, cancer, and stromal gene markers are highlighted. **b,** Cell type enrichment across compartments, expressed as observed-to-expected ratios (Ro/e). **c,** Mutational signature enrichment in CC samples. **d,** Co-occurrence of mutational signatures in CC samples. **e,** Promoter methylation scores for APOBEC genes across cancer types. Wilcoxon rank-sum test, ****p<0.0001, *0.01≤p<0.05. **f,** Mean APOBEC expression in epithelial cells per sample across cancer types in the Visium HD dataset. Wilcoxon rank-sum test, ****p<0.0001, ***p<0.001, **p<0.01. **g,** Overlap of APOBEC mutational signatures and HPV infection status in CC samples.

**Extended Data Fig. 3:**
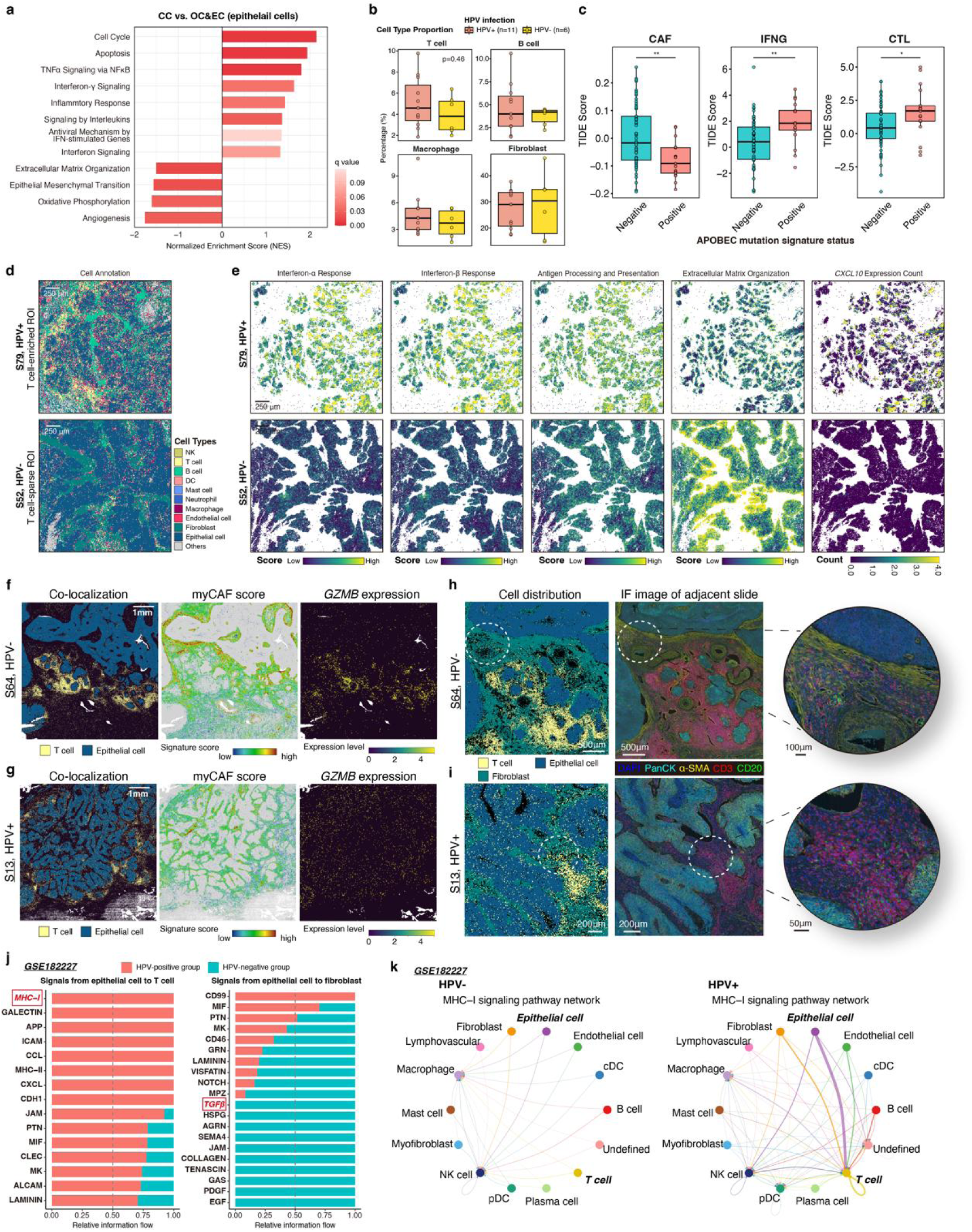
Spatial differences in CC by HPV infection status. **a,** GSEA for epithelial cells in CC versus non-CC samples in the Visium HD dataset. FDR-adjusted permutation test (10,000 permutations). **b,** Cell-type proportions stratified by HPV status in CC samples. Wilcoxon rank-sum test. **c,** CAF, IFN-γ, and CTL signature scores stratified by APOBEC mutational signature status in CC samples. Wilcoxon rank-sum test, **p<0.01, *p<0.05. **d,** Spatial cell-type maps (ROIs) from HPV+ (S79) and HPV-(S52) samples. **e,** Spatial maps of immune-related pathways in epithelial cells from ROIs in (**d**). **f,g,** Spatial maps of T-epithelial co-localization, myCAF scores, and *GZMB* expression in HPV-sample S64 (**f**) and HPV+ sample S13 (**g**) in the CC cohort. **h,i,** T-epithelial-myofibroblast interactions from Visium HD and paired IF images in HPV-sample S64 (**h**) and HPV+ sample S13 (**i**) in the CC cohort. **j,** Interaction strength of epithelial-T cell (left) and epithelial-fibroblast (right) pairs by HPV status in public scRNA-seq datasets. **k,** HPV-dependent modulation of MHC-I signaling interactions in public scRNA-seq.

**Extended Data Fig. 4:**
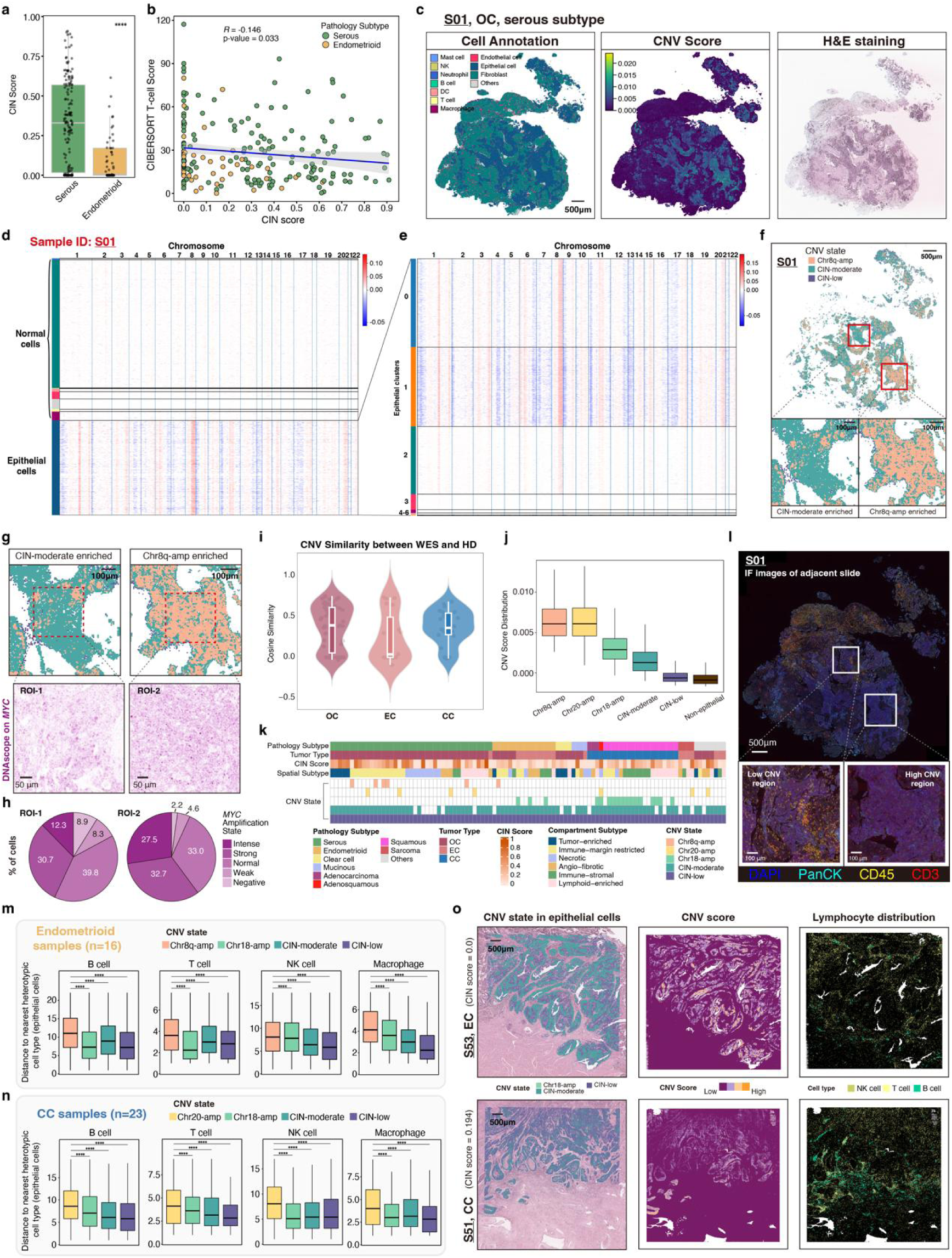
Spatial CNV profiling and cellular interactions in the FGT samples. **a,** CIN score distributions in serous vs. endometrioid tumors from WES data. Wilcoxon rank-sum test, ****p<0.0001. **b,** Correlation between CIN score and CIBERSORT T-cell score across serous and endometrioid subtypes using matched WES and RNA-seq data. Pearson’s R, two-sided t-test. **c,** Spatial mapping of cell-type annotation (left), CNV scores (middle), and H&E image for sample S01. CNV scores inferred from Visium HD data. **d,e,** Inferred CNV profiles for all cell types (**d**) and epithelial cells (**e**) in sample S01. **f,** Spatial distribution of three CNV states in sample S01, with representative epithelial ROIs. **g,** Representative ROIs correspond to distinct subclone regions in (**f**) from sample S01, along with spatial visualization of DNA probes for MYC (purple) of these ROIs from adjacent slide of S01. **h,** Percentages of cells exhibiting distinct *MYC* amplification states within the two ROIs. Cell counts and MYC probe signals were quantified using HALO software. **i,** Cosine similarity of Visium HD-derived CNV profiles with WES-derived CNV profiles across three cancer types. **j,** CNV score distributions across six CNV-defined cell states. **k,** Frequency of CNV states across 100 samples. **l,** Validation for T cells (CD45+ CD3+) exclusion from high CNV region using IF in adjacent slide. **m,n,** Spatial distance from epithelial cells to nearest other cell types in endometrioid (**m**) and CC (**n**) samples, stratified by CNV state. Wilcoxon rank-sum test, ****p<0.0001. **o,** Spatial characterization of CNV states, CNV scores, and immune infiltration in representative sections from EC (top) and CC (bottom) samples.

**Extended Data Fig. 5:**
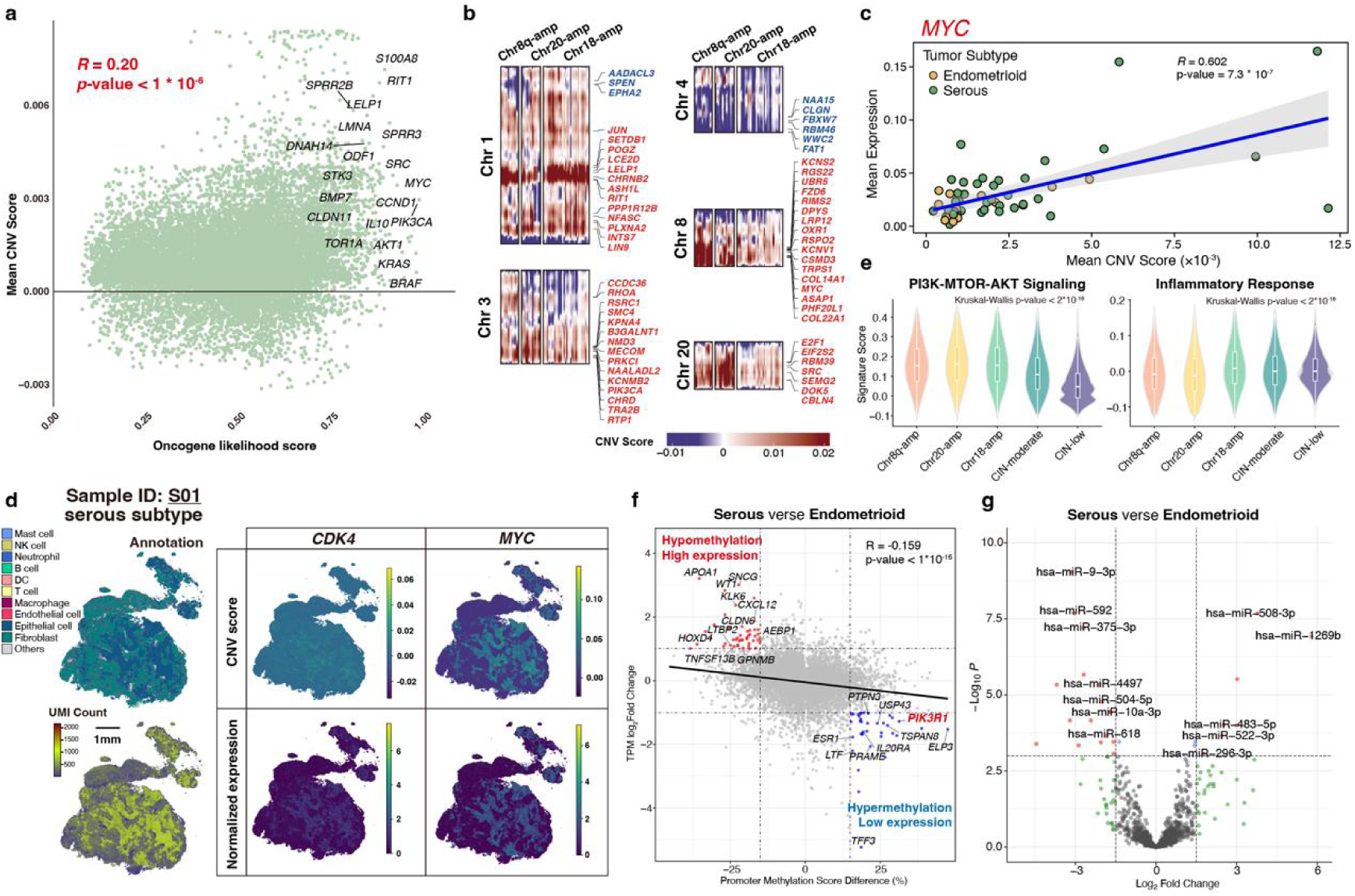
Spatial characterization of CNV states and oncogenic programs. **a,** Scatterplot showing correlation between mean Visium HD-derived CNV score and oncogene likelihood scores. Pearson’s R. Each dot represents a single gene. **b,** Chromosomal CNV profiles stratified by CNV state (Visium HD-derived), annotated for oncogenes and tumor suppressors. **c,** Correlation between mean CNV score with *MYC* expression across serous and endometrioid tumors (Visium HD dataset). Each dot represents one sample. Pearson’s R. **d,** Spatial characterization of cell annotations (top left), UMI counts (bottom left), Visium HD-derived CNV scores (top right), and *CDK4/MYC expression (bottom right)* from sample S01. **e,** Violin plots showing cancer-related pathway activity across CNV states. Kruskal-Wallis test. **f,** Integrative analysis of differentially methylated promoters and differentially expressed genes between serous and endometrioid subtypes. Each dot represents one gene. **g,** Differential miRNA expression between serous and endometrioid subtypes. labeled miRNAs with criteria |log₂ FC| > 1.5, adj.p < 0.001, mean count > 20.

**Extended Data Fig. 6:**
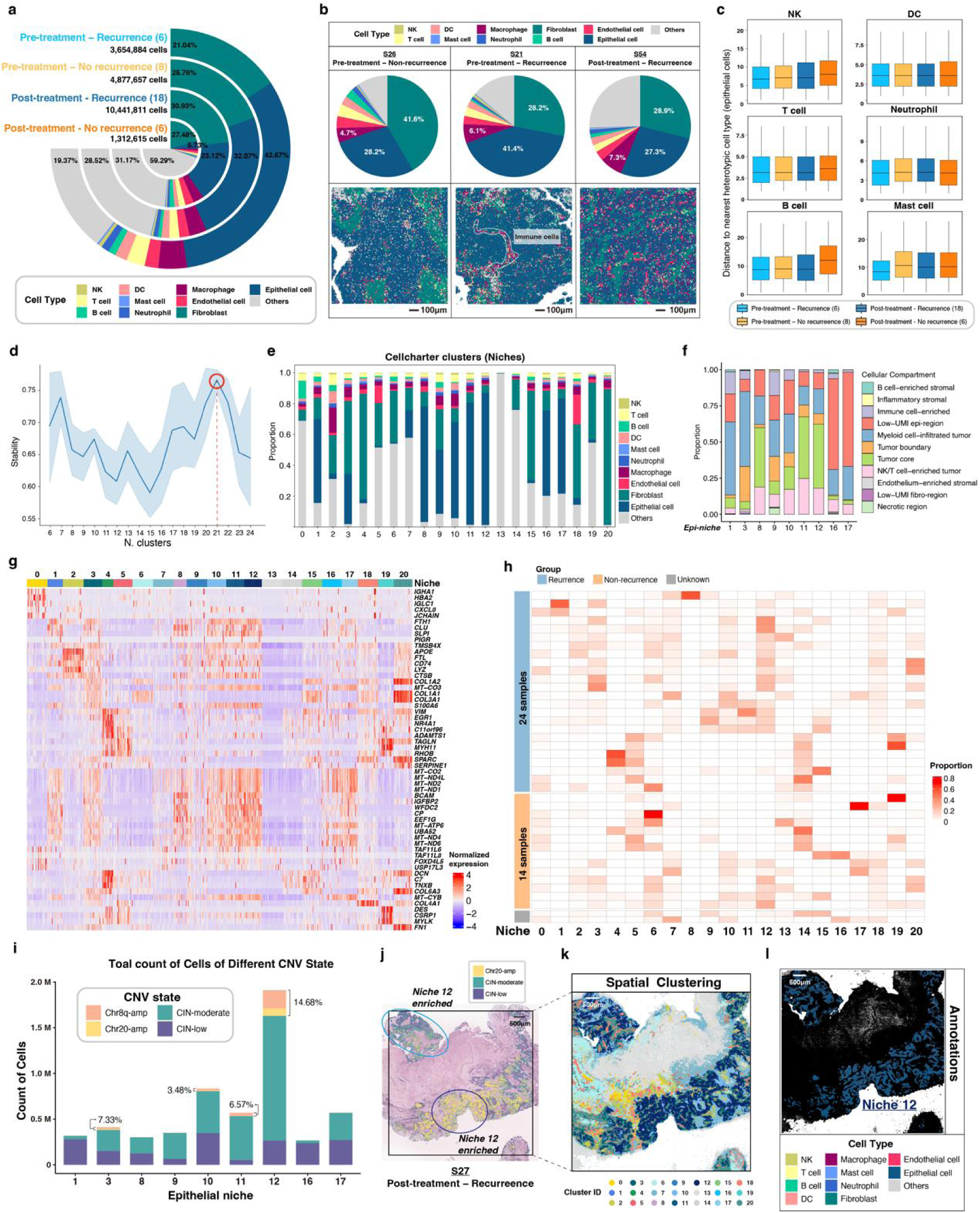
Spatial and cellular characterization of HGSOC samples across treatment and recurrence groups. **a,** Circular bar chart showing fibroblast, epithelial cell, and other cell-type proportions by treatment and recurrence group. **b,** Cell-type composition and spatial distribution of macrophages in representative ROIs across treatment and recurrence groups. **c,** Distances from epithelial cells to specific neighboring cell types, by treatment and recurrence groups. **d,** Cluster stability across different cluster numbers (k). **e,** Cell-type distributions per spatial cluster. **f,** Stacked bars of epithelial cells across cellular compartments. **g,** Heatmap of top differentially expressed genes per niche. **h,** Heatmap of niche enrichment across samples by recurrence status. **i,** Stacked bar plots of epithelial cell counts per niche with CNV states labeled. **j-l,** Spatial characterization of CNV states **(j)**, cluster annotations **(k)**, and cell-type annotations **(l)** in sample S27.

**Extended Data Fig. 7:**
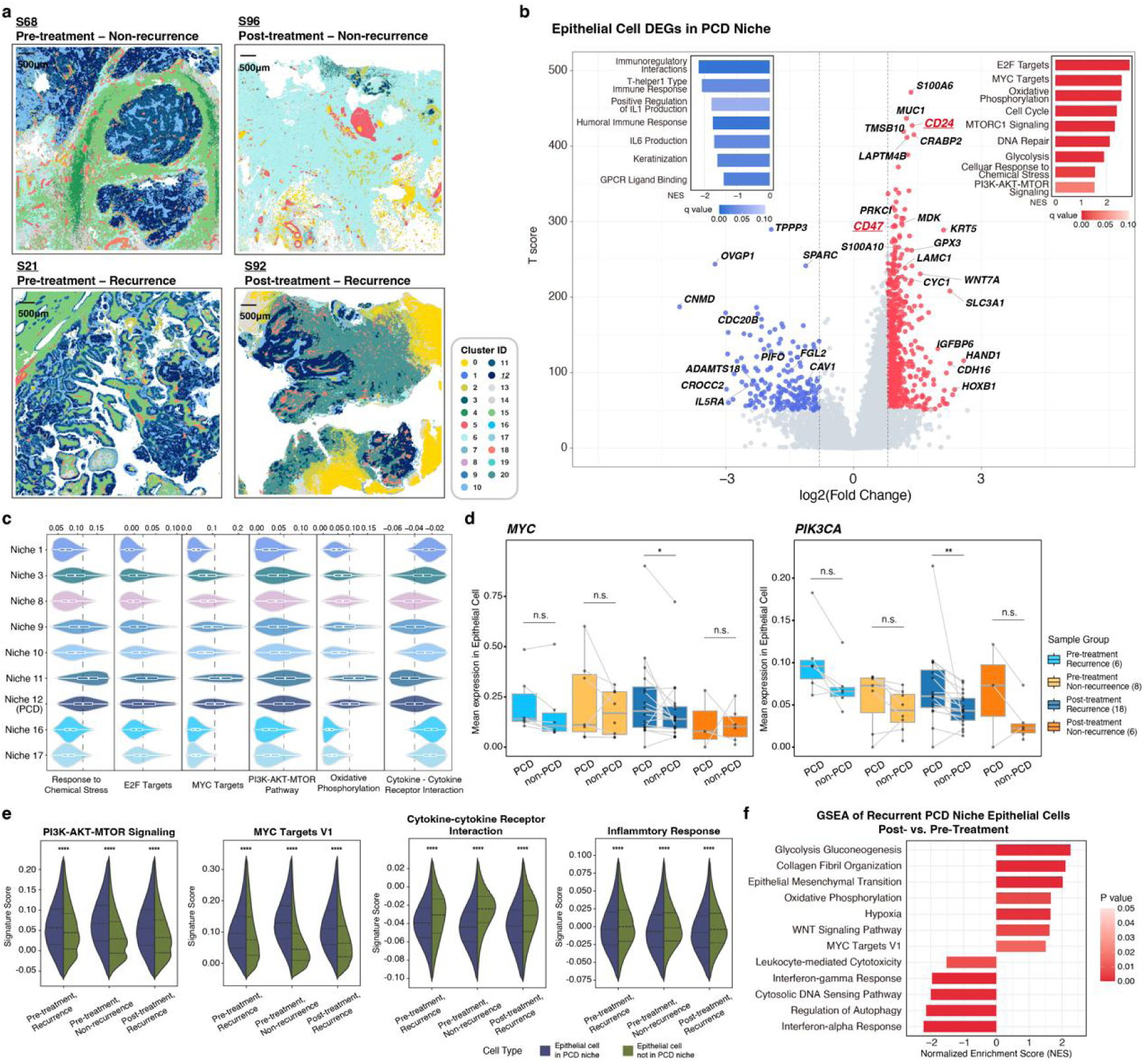
Pathways and cell niches linked to treatment and recurrence in HGSOC samples. **a,** Spatial clustering of Proliferating Core Desert (PCD) niches in representative HGSOC samples across treatment and recurrence groups. **b,** DEG analysis in PCD versus non-PCD epithelial niches with pathway enrichment of up-(top right) and downregulated (top left) genes. **c,** Violin plots showing cancer-related pathway levels across epithelial niches. **d,** CNV-driven oncogene (*MYC*, *PIK3CA*) expression in PCD versus non-PCD niches. Wilcoxon signed-rank test, **p<0.01, *0.01≤p<0.05, n.s. non-significant. **e,** Violin plots showing cancer-related pathway levels in epithelial cells inside versus outside PCD niche, stratified by treatment and recurrence groups. **f,** GSEA of epithelial cells in PCD niche from recurrent HGSOC samples, post-treatment versus pre-treatment.

**Extended Data Fig. 8:**
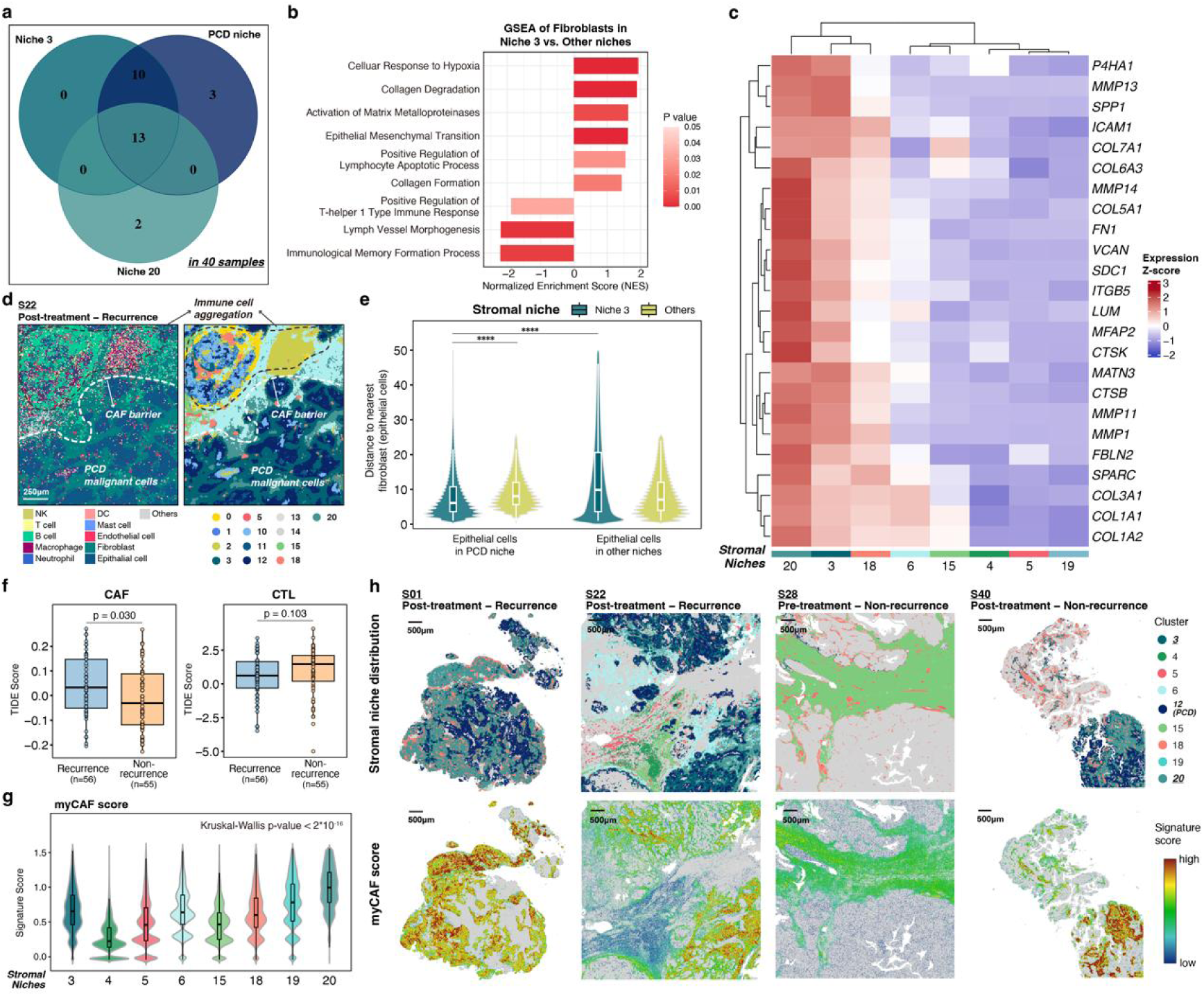
CAF-associated pathways and spatial interactions with PCD niche. **a,** Venn diagram of co-occurrence of niche 3, niche 20, and PCD niche across 40 HGSOC samples. **b,** GSEA results in niche 3 versus other fibroblasts. **c,** Heatmap of ECM remodeling gene expression in fibroblasts from stromal niches. **d,** Spatial co-localization of epithelial cells, CAFs, and immune cells (left), with niche annotations assigned to the observed cellular distributions (right), in the ROI from a recurrent sample (S22). **e,** Spatial distance from epithelial cells to nearest fibroblasts, stratified by epithelial cell and fibroblast location. Wilcoxon rank-sum test, ****p<0.0001. **f,** Distribution of CAF and CTL signature scores, by recurrence status in HGSOC samples. Wilcoxon rank-sum test. **g,** Violin plots showing myCAF signature scores in fibroblasts across stromal niches. Kruskal-Wallis test. **h,** Spatial mapping of stromal niches (top) and myCAFs (bottom) in four representative samples (S01, S22, S28, S40), by treatment and recurrence status.

**Extended Data Fig. 9:**
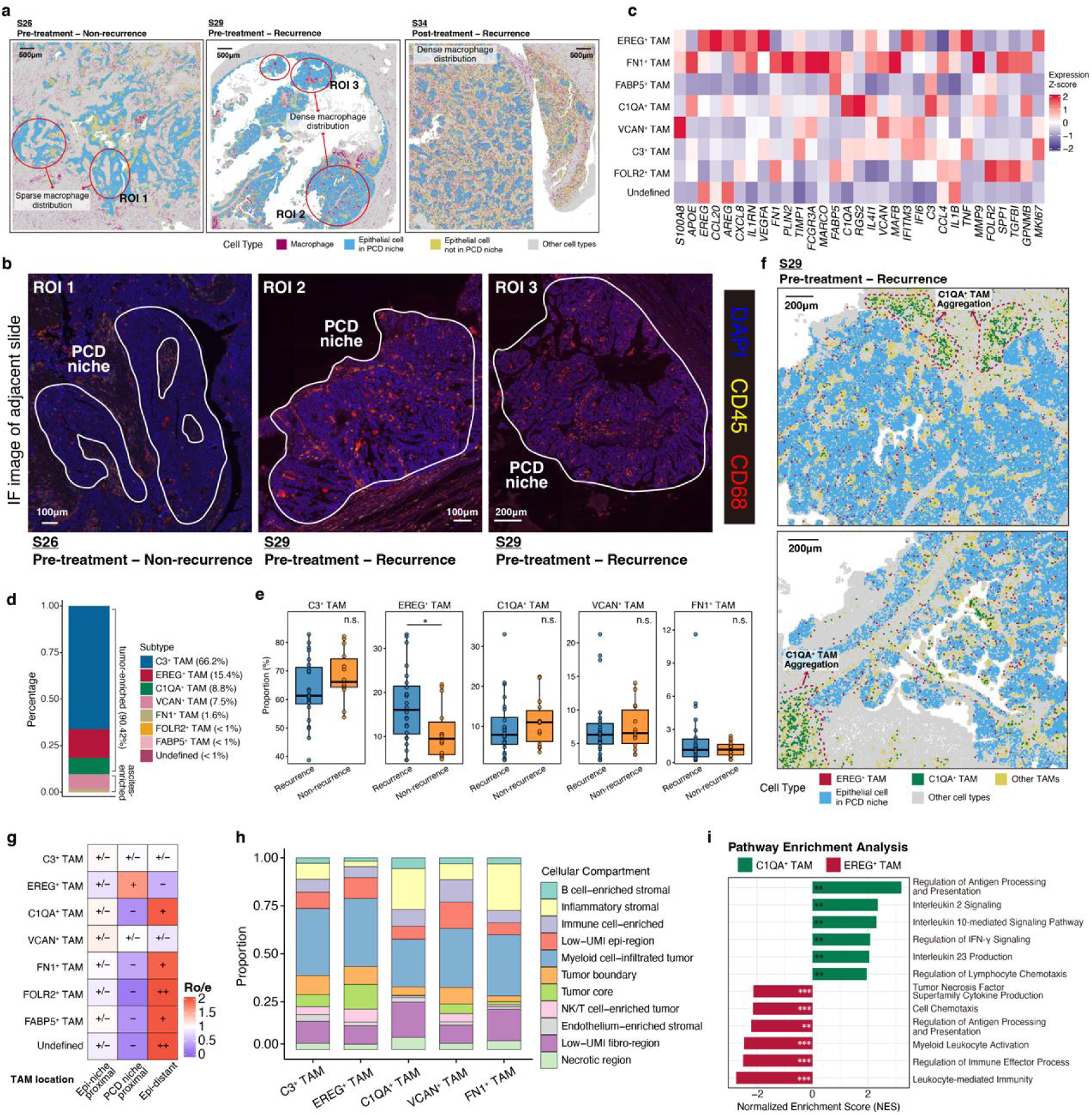
Tumor-associated macrophages (TAMs) in proximity to PCD niche. **a,b,** Spatial distribution of TAMs in three representative samples (S26, S29, S34) (**a**) and IF images of corresponding ROIs (**b**). **c,** Heatmap of TAM subtype marker expression from Visium HD annotations across single cell-based annotated TAMs in HGSOC from the dataset. **d.** Stacked bar plot showing the proportions of TAM subtypes in HGSOC samples (Visium HD dataset). **e,** Distribution of TAM subtype proportions by recurrence status in HGSOC samples. Wilcoxon rank-sum test, *0.01≤p<0.05. **f,** Spatial maps of EREG+ TAMs, C1QA+ TAMs, and other TAMs in two ROIs from S29, highlighting C1QA+ TAM aggregation outside PCD niche. **g,** Spatial preference of TAM subtypes, summarized as Ro/e ratios. **h,** Stacked bars of TAM proportions across cellular compartments, ordered by TAM subtype. **i,** GSEA of pathways upregulated in C1QA+ TAMs and downregulated in EREG^+^ TAMs.

**Extended Data Fig. 10:**
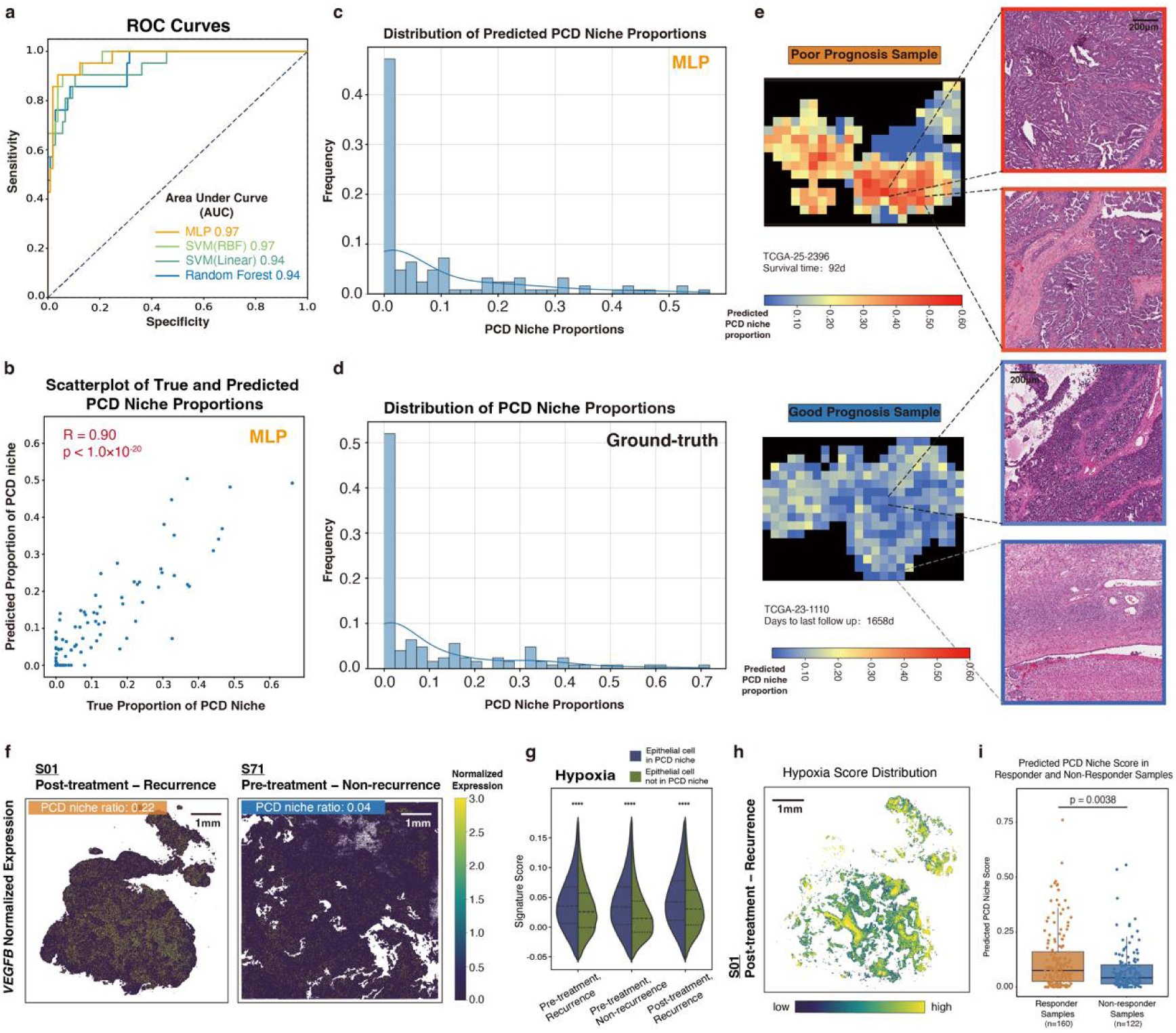
Machine learning performance and spatial features of PCD niches. **a,** Performance of selected machine learning models, evaluated by AUC on the test set without additional hyperparameter tuning. **b,** Scatterplot comparing observed (x-axis) versus predicted (y-axis) PCD niche proportions in the test set (n=126). Pearson’s R. **c,d,** Distributions of predicted (**c**) and true (**d**) PCD niche proportions across validation set (n=125). **e,** Cropped H&E patches from a poor-prognosis, PCD-high sample (top) and a good-prognosis, PCD-low sample (bottom). **f,** Spatial characterization of *VEGFB* expression in a PCD-enriched (left) and PCD-deficient (right) sample. **g,** Violin plot showing hypoxia pathway levels in epithelial cells inside versus outside PCD, stratified by treatment and recurrence groups. **h,** Spatial characterization of hypoxia signature in epithelial cells from S01. **i,** Predicted PCD niche proportions in bevacizumab-responder versus non-responders.. Wilcoxon rank-sum test.

